# Infected macrophages engage alveolar epithelium to metabolically reprogram myeloid cells and promote antibacterial inflammation

**DOI:** 10.1101/2020.02.29.970004

**Authors:** Xin Liu, Mark A. Boyer, Alicia M. Holmgren, Sunny Shin

## Abstract

Alveolar macrophages are the primary immune cells that first detect lung infection. However, only one macrophage patrols every three alveoli. How this limited number of macrophages provides protection is unclear, as numerous pathogens block cell-intrinsic immune responses. The intracellular pathogen *Legionella pneumophila* inhibits host translation, thereby impairing the ability of infected macrophages to produce critical cytokines. Nevertheless, infected macrophages induce an IL-1-dependent inflammatory response by recruited myeloid cells that controls infection. Here, we show that collaboration with the alveolar epithelium is critical, in that IL-1 instructs the alveolar epithelium to produce GM-CSF. Intriguingly, GM-CSF drives maximal cytokine production in bystander myeloid cells by enhancing PRR-induced glycolysis. Our findings reveal that alveolar macrophages engage alveolar epithelial signals to metabolically reprogram myeloid cells and amplify antibacterial inflammation.

**One Sentence Summary:** The alveolar epithelium is a central signal relay between infected and bystander myeloid cells that orchestrates antibacterial defense.

## Main Text

The lung is one of the largest mucosal surfaces in the body. Alveolar macrophages are the primary immune cells present in the distal lung and among the first to encounter inhaled bacterial pathogens. Antimicrobial immunity is thought to be initiated upon sensing of pathogen-associated molecular patterns (PAMPs) by pattern recognition receptors (PRRs). Subsequently, there is recruitment of bystander myeloid cells, such as monocytes and neutrophils, which amplify inflammatory responses important for controlling infection (*1, 2*). However, only one alveolar macrophage patrol every three alveoli (*3, 4*), and it is unclear how a limited number of macrophages can patrol such a vast surface area. In addition, numerous pathogens disarm infected macrophages by deploying virulence factors that impair cell-intrinsic PRR signaling. Thus, how an effective immune response can be generated against pathogens remains a fundamental question, and suggest the existence of additional mechanisms that promote antimicrobial defense.

To define how antibacterial immunity is generated in response to pathogens that interfere with immune signaling, we utilized *Legionella pneumophila*, which causes the severe pneumonia Legionnaires’ disease (*5, 6*). *Legionella* infects and replicates within alveolar macrophages (*7*). To do so, *Legionella* employs a bacterial type IV secretion system (T4SS) to translocate effector proteins into the host cell (*8-10*). These effectors manipulate numerous host processes to promote bacterial replication within macrophages (*8, 10, 11*). A subset of these effectors potently inhibit host translation (*12-17*). As a result, infected macrophages are incapable of making cytokines, such as TNF and IL-12, which are required to control infection (*18*). However, infected cells still translate and release IL-1α and IL-1β (*18-20*). Critically, IL-1R signaling elicits TNF and IL-12 production by bystander myeloid cells in the lung (*18*). These findings indicate that IL-1 released by *Legionella*-infected macrophages drives the induction of critical cytokines by bystander cells, allowing for immune bypass of the bacterial block in host translation to enable control of infection.

IL-1R signaling is important in control of many microbial infections, but how IL-1 instructs bystander cytokine responses is poorly understood. IL-1R is expressed by both hematopoietic and non-hematopoietic cells (*21-23*). Whether cell-intrinsic IL-1R signaling instructs myeloid cells to produce cytokines is unknown. Here, we report that IL-1R signaling on alveolar type II epithelial cells (AECII), rather than immune cells, was critical in driving cytokine production by bystander myeloid cells and control of *Legionella* infection. Mechanistically, IL-1 was both necessary and sufficient to instruct AECII to make the cytokine GM-CSF, which in turn was required for optimal production of IL-1, TNF, and IL-12 by myeloid cells in the infected lung. Intriguingly, GM-CSF promoted metabolic reprogramming of MCs to undergo heightened aerobic glycolysis, which was required for robust cytokine expression. Thus, the alveolar epithelium is a central signal relay between infected and bystander myeloid cells that optimizes inflammatory cytokine responses and host defense via GM-CSF-induced metabolic reprogramming.

## Results

### Non-hematopoietic IL-1R signaling controls inflammatory cytokine production by myeloid cells and antibacterial defense

To define how antibacterial immunity is generated in response to pathogens that interfere with immune signaling, we utilized *Legionella pneumophila*. Upon infecting alveolar macrophages, *Legionella* injects effectors that potently block host protein synthesis, impairing the ability of infected macrophages to produce inflammatory cytokines (*18*). Nevertheless, *Legionella*-infected macrophages still synthesize and release IL-1α and IL-1β (*18-20*). Our previous work revealed that IL-1R signaling was required for bystander MCs and other myeloid cells to produce TNF and IL-12 (*18*), which are required to control infection (*24, 25*).

IL-1R is expressed by both hematopoietic and stromal cells in the lung, but IL-1 stimulation of myeloid cells *in vitro* does not induce high expression of inflammatory cytokines (*18, 26*), raising the question of which cell types respond to IL-1. We generated bone marrow (BM) chimeras in which IL-1R expression was confined to the stromal compartment by providing *Il1r1*^*-/-*^ BM to lethally irradiated WT mice (*Il1r1*^*-/-*^ →WT) or the hematopoietic compartment by providing WT BM to lethally irradiated *Il1r1*^*-/-*^ mice (WT→*Il1r1*^*-/-*^), along with control chimeras (WT→WT or *Il1r1*^*-/-*^ →*Il1r1*^*-/-*^). Following hematopoietic reconstitution (Figure S1A), we intranasally infected the chimeras with a sublethal dose of *Legionella* and assessed cytokine production in the bronchoalveolar lavage fluid (BAL) 24 hours post-infection. Surprisingly, mice lacking stromal IL-1R expression had a significant defect in TNF, IL-6, and IL-12 levels compared with WT→WT control mice or *Il1r1*^*-/-*^ →WT mice (Figure 1A).

**Fig. 1.**
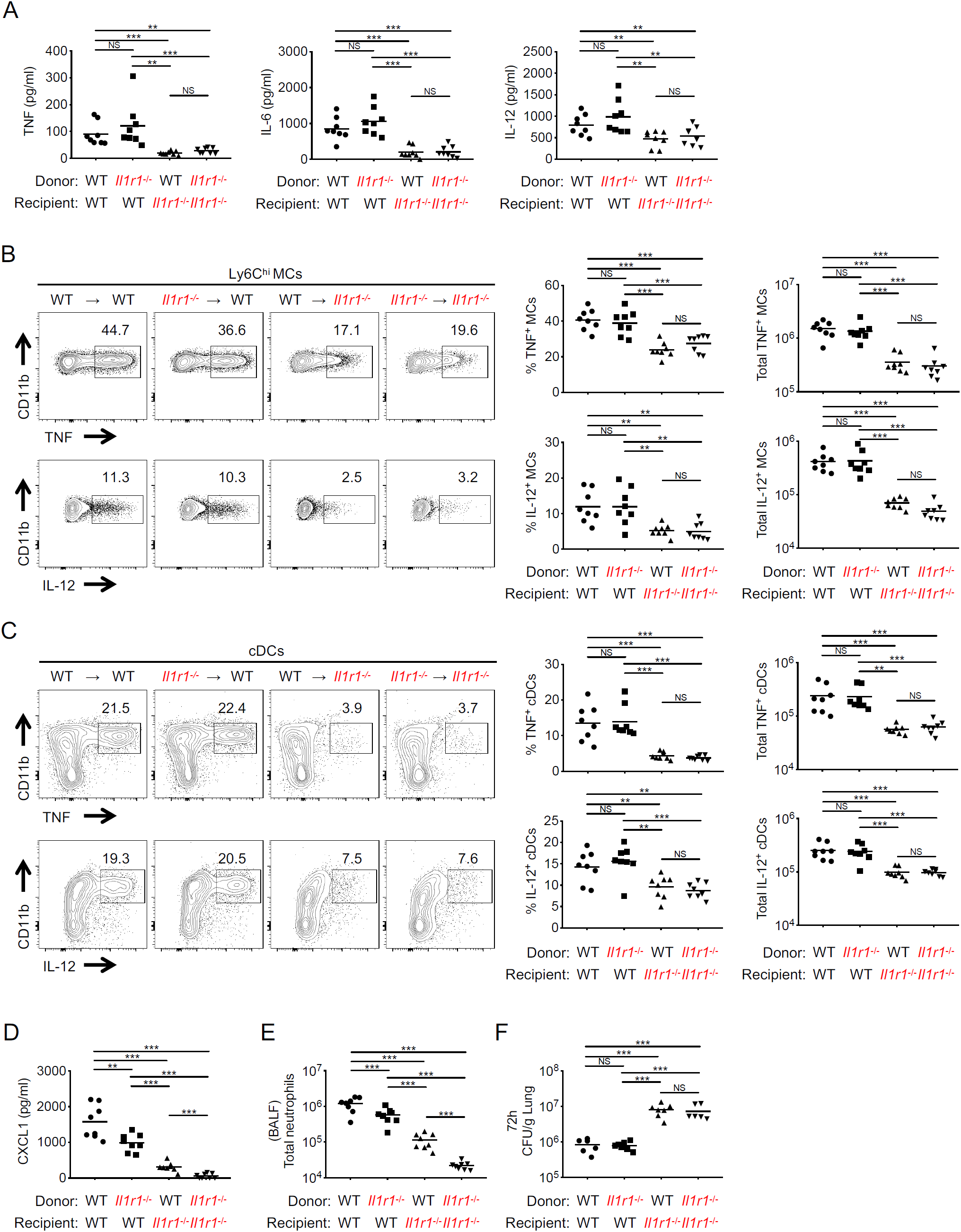
IL-1R signaling in nonhematopoietic cells orchestrates antibacterial defense. Indicated BM chimeras were intranasally infected with *Legionella (L*.*p*.). (A) TNF, IL-6, and IL-12 levels in the bronchoalveolar lavage fluid (BAL) at 24 hours post-infection (hpi). (B and C) Representative flow cytometric plots showing intracellular staining for TNF and IL-12 in Ly6C^hi^ MCs (B) and cDCs (C) from the lungs at 24 hpi. Frequency and total number of TNF- and IL-12-producing MCs (B) and cDCs (C) are shown. (D and E) CXCL1 levels (D) and total number of neutrophils (E) in the BAL at 24 hpi. (F) *L*.*p*. CFUs in the lungs of chimeric mice at 72 hpi. Data shown are the pooled results of two independent experiments with 3-4 mice per group in each experiment. NS, not significant; **p < 0.01; and ***p < 0.001 (one-way ANOVA with Turkey’s multiple comparisons test). See also Figure S1.

Ly6C^hi^ MCs and dendritic cells (DCs) are major producers of TNF and IL-12 during *Legionella* infection (*27, 28*). We observed that both the frequency and total number of TNF- or IL-12-producing Ly6C^hi^ MCs and DCs in the lung were significantly reduced in WT→*Il1r1*^*-/-*^ mice lacking stromal IL-1R expression (Figure 1B and 1C), although total numbers of lung MCs and cDCs in the different chimeras were equivalent (Figure S1B). Indeed, the observed cytokine defect in WT→*Il1r1*^*-/-*^ mice was indistinguishable from *Il1r1*^*-/-*^ →*Il1r1*^*-/-*^ mice. These data indicate that stromal IL-1R signaling was required for myeloid cells to produce inflammatory cytokines during infection. In contrast, both stromal and hematopoietic IL-1R signaling was necessary for optimal production of the chemokine CXCL1 and neutrophil recruitment, although loss of stromal IL-1R signaling had a greater impact (Figure 1D and 1E). Importantly, at this early timepoint, bacterial loads were equivalent in all chimeras (Figure S1C), indicating that decreased cytokine production in mice lacking stromal IL-1R signaling was not due to elevated levels of bacteria inhibiting protein synthesis.

Critically, loss of stromal IL-1R expression was associated with failure to control bacterial burden, as WT→*Il1r1*^*-/-*^ and *Il1r1*^*-/-*^ →*Il1r1*^*-/-*^ mice had significantly higher bacterial loads compared to WT→WT and *Il1r1*^*-/-*^ →WT mice at 72 hours post-infection (Figure 1F). To address whether there was a myeloid cell-intrinsic requirement for IL-1R signaling in cytokine production that is masked by stromal IL-1R signaling, we generated mixed BM chimeras in which lethally irradiated WT or *Il1r1*^*-/-*^ recipients were given a 1:1 ratio of *Il1r1*^*-/-*^ and WT BM. We observed equivalent frequencies of TNF- or IL-12-producing *Il1r1*^*-/-*^ or WT MCs and cDCs within the chimeric mice (Figure S1D and S1E), indicating that myeloid cell-intrinsic IL-1R signaling is not required. Thus, IL-1R signaling within stromal cells, but not hematopoietic cells, is required for myeloid cells to produce inflammatory cytokines and control of infection.

### IL-1R on alveolar type II epithelial cells is essential for cytokine production by myeloid cells and antibacterial defense

Alveolar type II epithelial cells (AECII) constitute 60% of alveolar epithelial cells (*29*), and can produce chemokines and cytokines in response to IL-1 (*30, 31*). To ask whether IL-1R specifically on AECII is required for myeloid cells to produce cytokines and antibacterial defense, we generated mice lacking IL-1R on AECII by crossing *Il1r1*^*fl/fl*^ mice with mice expressing CreER^T2^ under the control of the AECII-specific *Sftpc* promoter. Tamoxifen (TXF) treatment resulted in ∼80% deletion efficiency of the *Il1r1* locus in AECII from *Il1r1*^*fl/fl*^ *;Spc-cre*^*ERT2*^ mice compared to *Il1r1*^*fl/fl*^ mice (Figure 2A). Following infection, TXF-treated *Il1r1*^*fl/fl*^ *;Spc-cre*^*ERT2*^ mice produced significantly less TNF, IL-12, and IL-6 (Figure 2B), and had a log-increase in lung bacterial loads compared to *Il1r1*^*fl/fl*^ mice, whereas in the absence of TXF treatment, *Il1r1*^*fl/fl*^ *;Spc-cre*^*ERT2*^ and *Il1r1*^*fl/fl*^ mice had similar bacterial loads (Figure 2C). In contrast, and consistent with our data that hematopoietic IL-1R signaling is dispensable for controlling infection, we saw no difference in cytokine levels or bacterial loads between *Il1r1*^*fl/fl*^ and *Il1r1*^*fl/fl*^ *;Cd11c-cre* mice, which lack IL-1R expression on CD11c-expressing DCs and alveolar macrophages (Figure S2A and S2B). Finally, to test whether IL-1R expression on AECII is sufficient for inflammatory cytokine responses and control of infection, we limited IL-1R expression only to AECII by crossing mice containing a loxP-flanked disruptive sequence in the *Il1r1* gene and an IRES-tdTomato reporter (*Il1r1*^*r/r*^) to *Spc-cre*^*ERT2*^ mice. TXF treatment would be expected to restore IL-1R expression in AECII in *Il1r1*^*r/r*^ *;Spc-cre*^*ERT2*^ mice lacking IL-1R in all other tissues. Following TXF treatment, approximately 60% of AECII were tdTomato-positive; by contrast, a small percentage of other stromal lung cells, including AECI and bronchial alveolar stem cell (BASC), were tdTomato-positive, indicating that IL-1R expression was predominantly restored in AECII (Figure 2D), whereas AECII from TXF-treated control *Il1r1*^*r/r*^ mice did not express tdTomato. Strikingly, the *Il1r1*^*r/r*^ *;Spc-cre*^*ERT2*^ mice exhibited complete restoration of TNF, IL-12, and IL-6 levels and decreased bacterial loads compared to control *Il1r1*^*r/r*^ mice (Figure 2E and 2F). Altogether, IL-1R signaling in AECII is both necessary and sufficient for bystander cytokine production and bacterial clearance.

**Fig. 2.**
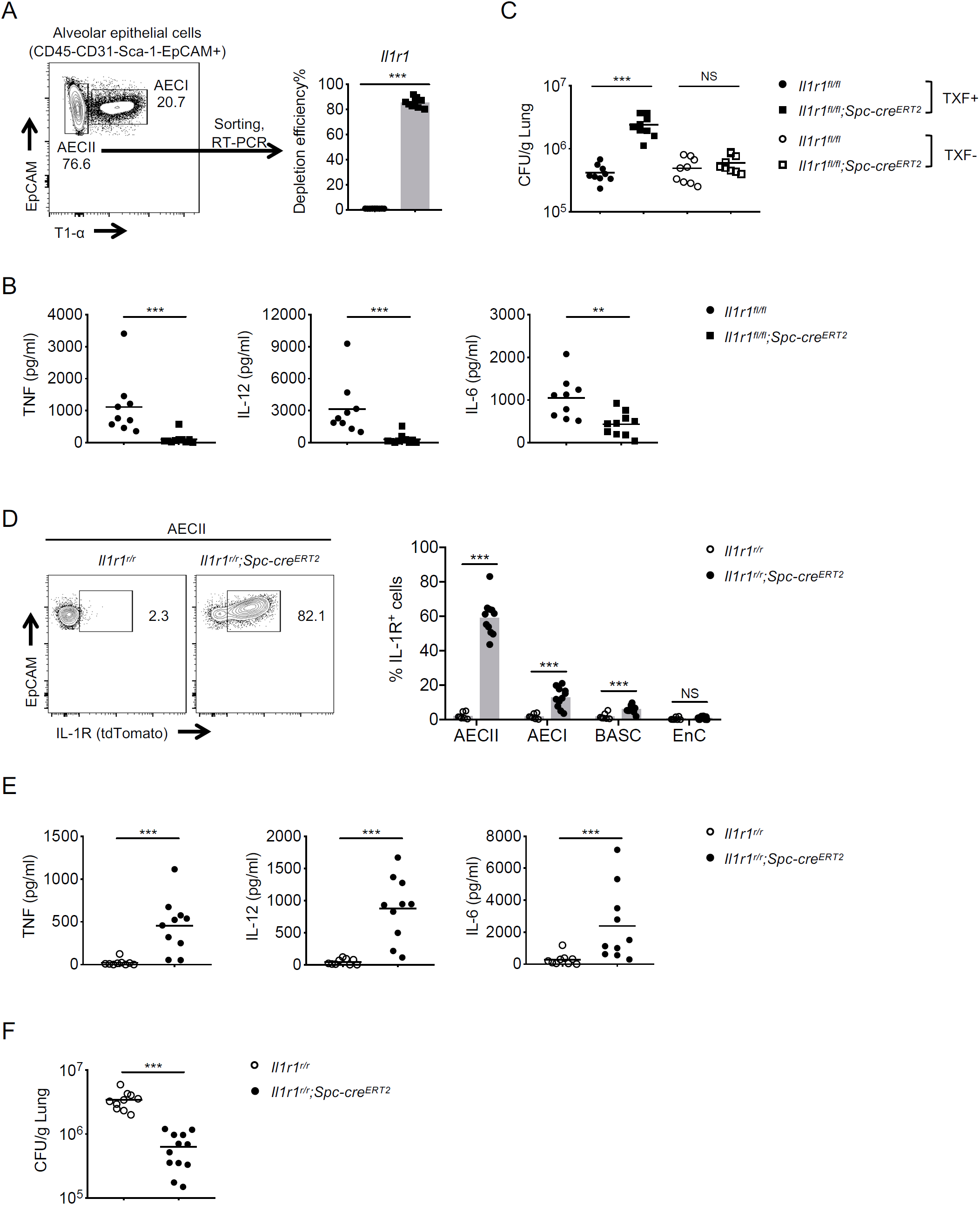
IL-1R expression on alveolar type II epithelial cells (AECII) is necessary and sufficient for antibacterial defense. (A) Representative flow cytometric plot showing the gating strategy for AECII (CD45-CD31-Sca-1^-^ EpCAM^+^ T1α^-^). Graph shows *Il1r1* deletion efficiency in AECII from *Il1r1*^*fl/fl*^ *;Spc-cre*^*ERT2*^ mice (▪) relative to *Il1r1*^*fl/fl*^ littermates (•) following tamoxifen (TXF) injection. (B and C) TNF, IL-12 and IL-6 levels in the BAL of TXF-injected *Il1r1*^*fl/fl*^ *;Spc-cre*^*ERT2*^ and *Il1r1*^*fl/fl*^ mice at 24 hpi (B). *L*.*p*. CFUs in the lungs of mice ± TXF injection at 72 hpi (C). (D) Representative flow cytometric plots showing tdTomato fluorescence as a measure of *Il1r1* expression in AECII from TXF-injected *Il1r1*^*r/r*^ *;Spc-cre*^*ERT2*^ and *Il1r1*^*r/r*^ mice at 24 hpi. Graph depicts frequency of IL-1R-expressing AECII, alveolar type I epithelial cells (AECI), bronchial alveolar stem cells (BASC), and lung endothelial cells (EnC). (E and F) TNF, IL-12 and IL-6 levels in the BAL at 24 hpi (E) and *L*.*p*. CFUs in the lungs at 72 hpi (F) of TXF-injected *Il1r1*^*r/r*^ *;Spc-cre*^*ERT2*^ and *Il1r1*^*r/r*^ mice. Data shown are the pooled results of three independent experiments with 3-4 mice per group in each experiment. NS, not significant; **p < 0.01; and ***p < 0.001 (unpaired t test). See also Figure S2.

### IL-1R signaling in AECII drives production of GM-CSF

We hypothesized that IL-1R signaling in AECII induces production of a soluble mediator that licenses myeloid cells to produce inflammatory cytokines. We therefore analyzed the BAL of WT and *Il1r*^*-/-*^ mice 24 hours post-infection using a Luminex assay to identify soluble factors that required IL-1R signaling for their induction. Many cytokines, including GM-CSF, were reduced in the BAL of *Il1r*^*-/-*^ mice compared to WT mice (Figure S2C). Intriguingly, AECII can produce GM-CSF (*31, 32*), and GM-CSF regulates alveolar macrophage differentiation (*33, 34*). Furthermore, GM-CSF is required for host defense against several lung pathogens, such as *Mycobacterium tuberculosis, Aspergillus fumigatus, Blastomyces dermatitidis*, and influenza virus (*35-38*). Following *Legionella* infection, GM-CSF was robustly induced in the lungs by 24 hours, and declined to background levels by 72 hours (Figure 3A). Critically, GM-CSF levels were nearly undetectable in *Il1a*^*-/-*^, *Il1b*^*-/-*^, or *Il1r1*^*-/-*^ mice at 24 hours post-infection (Figure 3B), indicating that IL-1R signaling induces GM-CSF during infection.

**Fig. 3.**
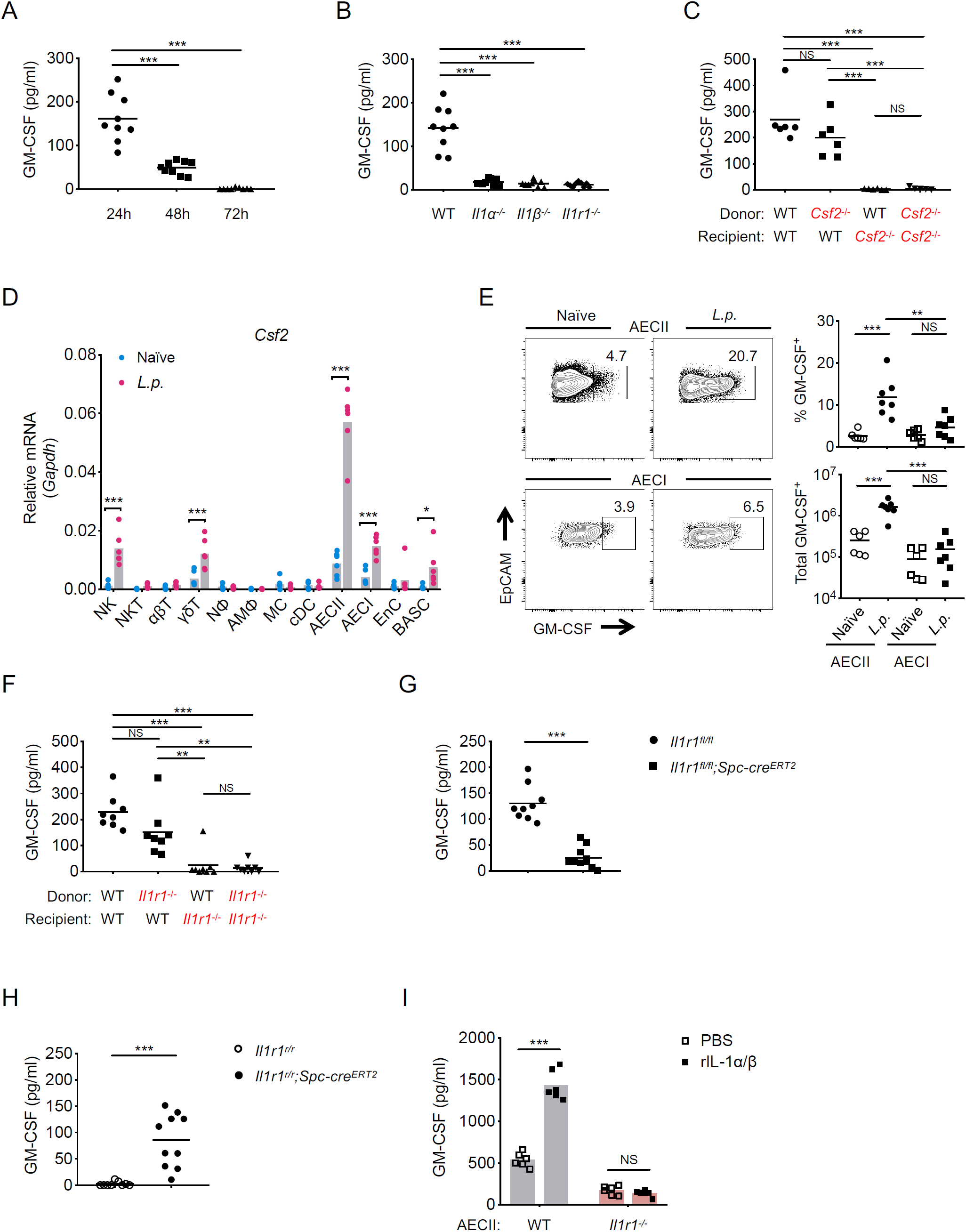
IL-1R signaling instructs AECII to produce GM-CSF during pulmonary *Legionella* infection. (A-C) GM-CSF levels in the BAL of C57BL/6 mice at 24, 48, and 72 hpi (A), WT, *Il1a*^*-/-*^, *Il1b*^*-/-*^ and *Il1r1*^*-/-*^ mice at 24 hpi (B), and WT:*Csf2*^*-/-*^ BM chimeras at 24 hpi (C). (D)*Csf2* transcript levels in the indicated cell types isolated from the lungs of naïve and *L*.*p*.-infected C57BL/6 mice at 24 hpi. (E) Representative flow cytometric plots showing intracellular staining for GM-CSF in AECII and AECI from the lungs of naïve and *L*.*p*.-infected C57BL/6 mice at 24 hpi. Frequency and total number of GM-CSF-producing AECII and AECI. (F-H) GM-CSF levels in the BAL of WT:*Il1r1*^*-/-*^ BM chimeras (F), *Il1r1*^*fl/fl*^ *;Spc-cre*^*ERT2*^ and *Il1r1*^*fl/fl*^ mice (G), and *Il1r1*^*r/r*^ *;Spc-cre*^*ERT2*^ and *Il1r1*^*r/r*^ mice (H) at 24 hpi. (I) GM-CSF levels in the supernatants of primary WT or *Il1r1*^*-/-*^ AECII treated with 10 ng/ml each of recombinant IL-1α and IL-1β (rIL-1α/β) or PBS vehicle control and infected with *L*.*p*. (MOI=5) at 12 hpi. Data shown are the pooled results of three (A, B, G, and H) or two (C-F) independent experiments with 3-4 mice per condition in each experiment. Data shown in I are the pooled results of two independent experiments with 3 mice and triplicate wells per condition in each experiment. NS, not significant; *p<0.05, **p<0.01; and ***p<0.001 (one-way ANOVA with Turkey’s multiple comparisons test for A-C and F; unpaired t test for D, E and G-I). See also Figure S2.

To determine whether a hematopoietic or stromal cell type produced GM-CSF during infection, we generated BM chimeras in which GM-CSF expression was confined to stromal cells by providing *Csf2*^*-/-*^ BM to lethally irradiated WT mice (*Csf2*^*-/-*^ →WT), or to hematopoietic cells by providing WT BM to lethally irradiated *Csf2*^*-/-*^ mice (WT→*Csf2*^*-/-*^), or appropriate control chimeras (WT→WT and *Csf2*^*-/-*^ →*Csf2*^*-/-*^). Mice with a WT stromal compartment (*Csf2*^*-/-*^ →WT) produced GM-CSF at levels similar to control WT→WT mice following infection, whereas GM-CSF was undetectable in mice lacking stromal GM-CSF expression (Figure 3C). Moreover, following infection, *Csf2* transcript levels were the highest in AECII relative to other lung cell types (Figure 3D), and the frequency and total number of GM-CSF-producing AECII significantly increased, whereas AECI were unchanged (Figure 3E). These data indicate that AECII are the primary producers of GM-CSF during *Legionella* infection.

We next addressed whether GM-CSF production specifically requires IL-1R signaling in AECII. Critically, WT→*Il1r1*^*-/-*^ BM chimeras lacking stromal IL-1R expression or *Il1r1*^*fl/fl*^ *;Spc-cre*^*ERT2*^ mice lacking IL-1R in AECII had a significant defect in GM-CSF production compared to control WT→WT chimeras or *Il1r1*^*fl/fl*^ mice, respectively, following *Legionella* infection (Figure 3F and 3G). Consistently, GM-CSF production was fully restored in *Il1r1*^*r/r*^ *;Spc-cre*^*ERT2*^ mice expressing IL-1R only on AECII compared to *Il1r1*^*r/r*^ mice lacking IL-1R signaling (Figure 3H). Furthermore, recombinant IL-1 (rIL-1α/β) induced GM-CSF in purified WT AECII but not in *Il1r1*^*-/-*^ AECII during *in vitro Legionella* infection (Figure 3I). These results show that IL-1R signaling in AECII is both necessary and sufficient for AECII to produce GM-CSF during *Legionella* infection.

### GM-CSF signaling is critical for local inflammatory cytokine production and bacterial clearance

We hypothesized that AECII production of GM-CSF helps drive bystander myeloid cells to produce cytokines important for bacterial clearance. In support of this model, *Csf2*^*-/-*^ mice had significant defects in production of IL-1β and TNF at 24 hours post-infection, and TNF, IL-12, and IFNγ at 48 hours post-infection (Figure 4A), and a log increase in bacterial loads at 72 hours post-infection compared to WT mice (Figure 4B). To address the possibility that these effects could be due to a lack of alveolar macrophages in *Csf2*^*-/-*^ mice due to the critical role for GM-CSF in alveolar macrophage development, we acutely blocked GM-CSF with neutralizing antibodies, as this treatment leaves the alveolar macrophage compartment intact (Figure S3A). Notably, GM-CSF blockade led to significantly decreased levels of IL-1β at 24 hour post-infection, and TNF, IL-12, and IFNγ at 48 hour post-infection (Figure 4C), and a log increase in bacterial loads at 72 hours post-infection compared to isotype control-treated mice (Figure 4D). The frequency and total number of IL-1α- or IL-1β-producing MCs and cDCs were significantly reduced as well (Figure 4E and 4F). Importantly, these effects of GM-CSF deficiency were independent of its growth factor properties, as numbers of neutrophils, MCs, and DCs in the lungs of anti-GM-CSF-treated mice were unaffected (Figure S3B). Thus, GM-CSF is required for optimal cytokine responses and bacterial clearance.

**Fig. 4.**
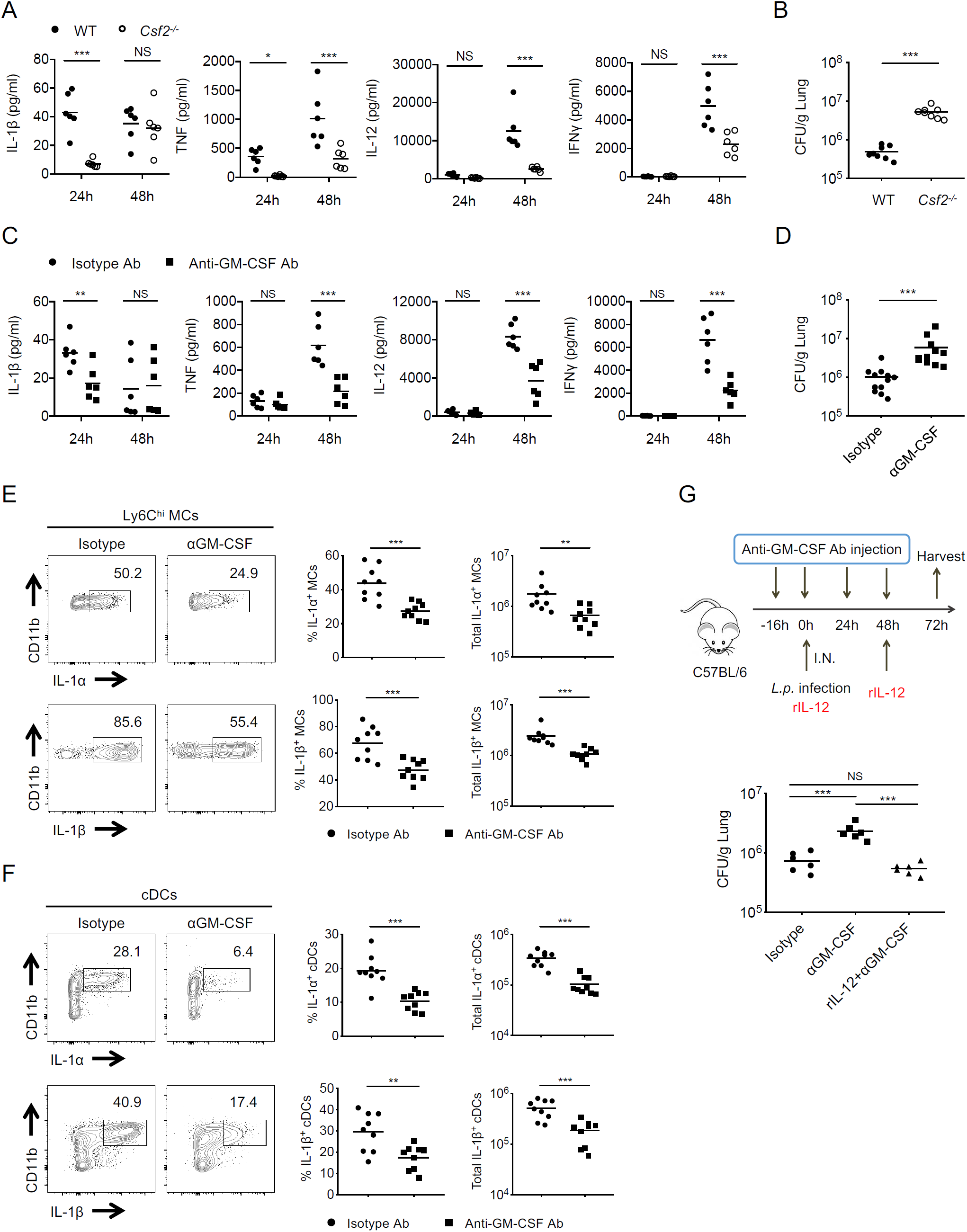
GM-CSF is required for maximal inflammatory cytokine production and bacterial clearance. (A and B) IL-1β, TNF, IL-12, and IFNγ levels in the BAL at 24 and 48 hpi (A) and *L*.*p*. CFUs in the lungs at 72 hpi (B) of WT and *Csf2*^*-/-*^ mice. (C and D) C57BL/6 mice injected with anti-GM-CSF or isotype control antibodies (Ab) prior to *L*.*p*. infection (see Method Details). IL-1β, TNF, IL-12, and IFNγ levels in the BAL at 24 hpi and 48 hpi (C). *L*.*p*. CFUs in the lungs at 72 hpi (D). (E and F) Representative flow cytometric plots showing intracellular staining for IL-1α and IL-1β in Ly6C^hi^ MCs (E) and cDCs (F) from the lungs of anti-GM-CSF or isotype control Ab-treated mice infected with *L*.*p*. at 24 hpi. Frequency and total number of IL-1α- and IL-1β-producing MCs (E) and cDCs (F) are shown. (G) *L*.*p*. CFUs in the lungs at 72 hpi of C57BL/6 mice intraperitoneally injected with anti-GM-CSF or isotype control Ab prior to *L*.*p*. infection or a third group injected with anti-GM-CSF Ab provided 500 ng recombinant IL-12p70 (rIL-12) intranasally at the time of infection and at day 2 post-infection (see Method Details). Data shown are the pooled results of two (A-C and G) or three (D-F) independent experiments or with 3-4 mice per condition for each experiment. NS, not significant; *p<0.05; **p < 0.01; and ***p < 0.001 (unpaired t test for A-F; one-way ANOVA with Turkey’s multiple comparisons test for G). See also Figure S3.

We next asked whether GM-CSF enhances cytokine production by myeloid cells distally within the systemic circulation or locally within the lung. We injected mice intravenously with anti-CD45-PE antibodies to label circulating immune cells, followed by staining of all lung cells to distinguish MCs within the vasculature from MCs that entered interstitial lung tissue. The number of interstitial and vascular MCs within the lungs of infected mice were significantly increased compared to naïve mice (Figure S3C). GM-CSF blockade of infected mice significantly decreased the frequency and total number of IL-1-producing interstitial and vascular MCs within the lung compared to isotype control-treated mice (Figure S3D). In contrast, there were substantially lower numbers of IL-1-producing peripheral blood MCs compared to lung MCs, with no difference in the numbers of IL-1-producing peripheral blood MCs in anti-GM-CSF- or isotype control-treated mice (Figure S3D). GM-CSF appears to have a potent local effect by licensing myeloid cells within the lung to produce cytokines during infection.

We next sought to determine whether GM-CSF controls *Legionella* infection primarily by inducing inflammatory cytokines or antimicrobial effector functions. As GM-CSF blockade led to a defect in IL-12 production (Figure 4C), and IL-12 is critical for control of *Legionella* (*24*), we tested the hypothesis that IL-12 was a downstream mediator of GM-CSF-dependent bacterial clearance. Administering recombinant IL-12 p70 to anti-GM-CSF-treated mice significantly reduced bacterial loads to levels similar to those in isotype control-treated mice (Figure 4G). Although GM-CSF treatment of macrophages can inhibit intracellular *M. tuberculosis* replication (*36*), there was no difference in *Legionella* replication within WT and *Csf2*^*-/-*^ BM-derived macrophages (BMDMs), or BMDMs treated with rGM-CSF (Figure S3E). GM-CSF can promote reactive oxygen species (ROS) production in neutrophils during *A. fumigatus* infection (*37*), but we observed no change in neutrophil ROS production in the lungs of anti-GM-CSF-treated mice, compared to isotype control-treated mice during *Legionella* infection (Figure S3F). Thus, AECII-derived GM-CSF promotes control of *Legionella* by inducing production of cytokines, such as IL-12, by myeloid cells.

### Myeloid cell-intrinsic GM-CSF signaling controls inflammatory cytokine production

GM-CSF could instruct myeloid cells to produce cytokines directly, or through yet another intermediary signal. We next asked whether myeloid cell-intrinsic GM-CSF signaling was required for cytokine production. We generated BM chimeric mice in which GM-CSF receptor expression was absent from hematopoietic cells by reconstituting lethally irradiated WT mice with *Csf2rb*^*-/-*^ BM, or mixed BM chimeras in which lethally irradiated WT mice were reconstituted with a 1:1 mixture of congenically marked WT and *Csf2rb*^*-/-*^ BM (Figure S4A). Notably, consistent with our previous findings with GM-CSF deficiency, there were significantly decreased IL-1α, IL-1β, IL-6, TNF, IL-12, and IFNγ levels in the BAL of *Csf2rb*^*-/-*^ →WT mice compared to WT→WT or mixed BM chimeric mice following *Legionella* infection (Figure 5A). Moreover, the frequency and total number of IL-1-producing MCs and cDCs were significantly reduced in *Csf2rb*^*-/-*^ →WT mice compared to WT→WT mice (Figure S4B and S4C), although there was no difference in total MC numbers and only a slight reduction in cDCs (Figure S4D). *Csf2rb*^*-/-*^ →WT mice also had a log increase in bacterial loads compared to WT→WT mice (Figure 5B), indicating a crucial role for hematopoietic GM-CSF receptor signaling in controlling infection.

**Fig. 5.**
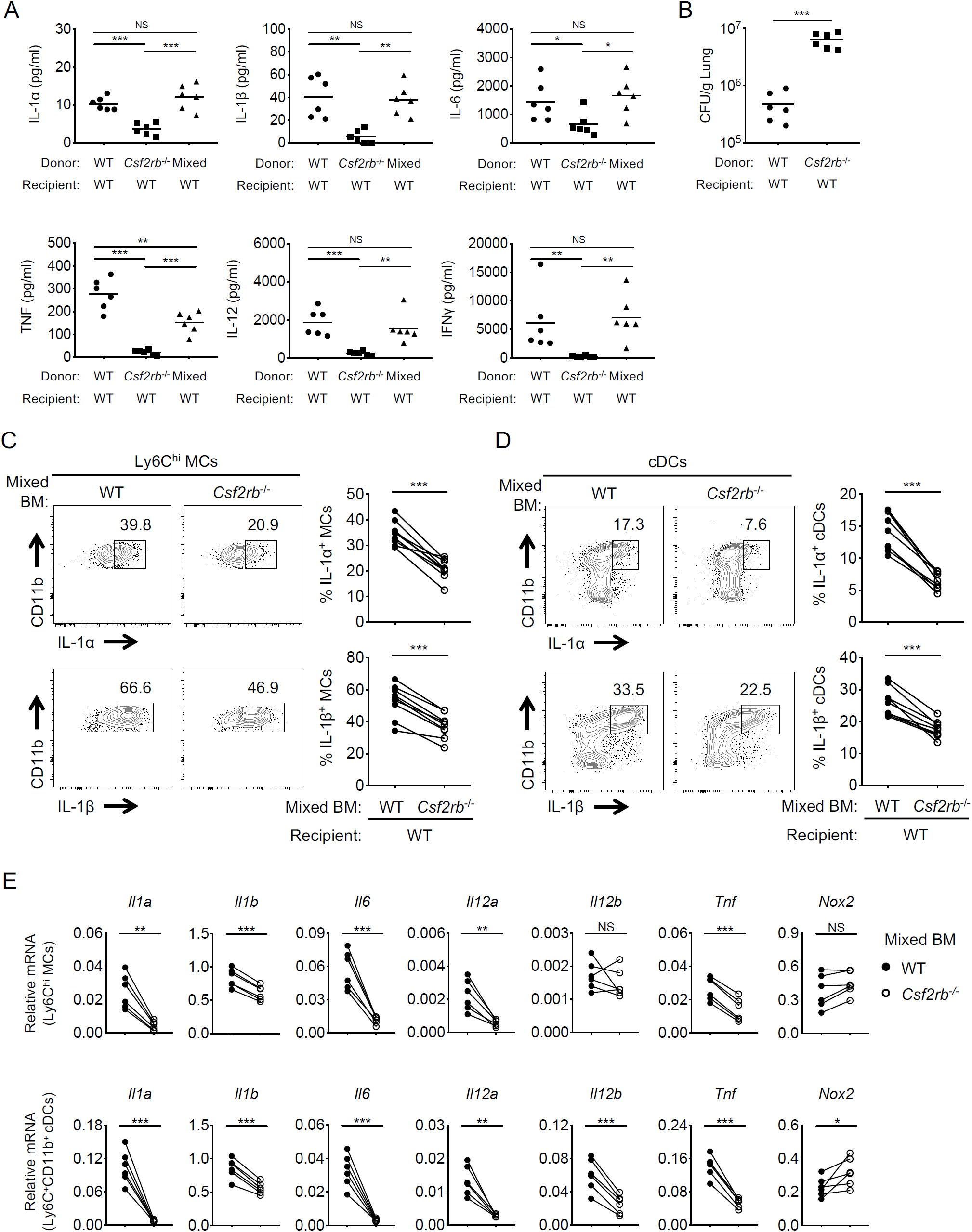
Cell-intrinsic GM-CSF receptor signaling is required for inflammatory cytokine production by myeloid cells. WT→WT, *Csf2rb*^*-/-*^ →WT, and 50% WT/50% *Csf2rb*^*-/-*^ →WT BM chimeras intranasally infected with *L*.*p*. (A) IL-1α, IL-1β, IL-6, TNF, and IL-12 levels in the BAL at 24 hpi and IFNγ at 48 hpi. (B) *L*.*p*. CFUs in the lungs of chimeric WT→WT and *Csf2rb*^*-/-*^ →WT mice at 72 hpi. (C and D) Representative flow cytometric plots and graphs depicting the frequency of IL-1α+ or IL-1β+ WT or *Csf2rb*^*-/-*^ MCs (C) and cDCs (D) from the lungs of 50% WT/50% *Csf2rb*^*-/-*^ →WT mixed BM chimeras at 24 hpi. Each line represents the paired values of WT and *Csf2rb*^*-/-*^ cells from a given mouse. (E) *Il1a, Il1b, Il6, Il12a, Il12b, Tnf*, and *Nox2* transcript levels in Ly6C^hi^ MCs and Ly6C^+^ CD11b^+^ cDCs isolated from the lungs of 50% WT/50% *Csf2rb*^*-/-*^ →WT mixed BM chimeras at 24 hpi, as quantified by qRT-PCR. Each line represents the paired values of WT and *Csf2rb*^*-/-*^ cells from a given mouse. Data shown are the pooled results of two (A, B, and E) or three (C and D) independent experiments with 3 mice per experiment. NS, not significant; *p < 0.05; **p < 0.01; and ***p < 0.001 (one-way ANOVA with Turkey’s multiple comparisons test for A; unpaired t test for B; Wilcoxon matched-pairs signed rank test for C-E). See also Figure S4.

Critically, in mixed BM chimeras, the presence of WT cells was unable to rescue cytokine production by *Csf2rb*^*-/-*^ cells, as the frequency of IL-1-producing *Csf2rb*^*-/-*^ MCs and cDCs was significantly reduced relative to WT cells within the same mouse (Figure 5C and 5D). Moreover, *ll1a, Il1b, Il6, Il12a*, and *Tnf* transcript levels were significantly reduced in *Csf2rb*^*-/-*^ Ly6C^hi^ MCs and Ly6C^+^ CD11b^+^ DCs, compared to WT cells within the same mouse (Figure 5E). Thus, myeloid cell-intrinsic GM-CSF signaling is critical for their ability to produce cytokines during *Legionella* infection.

### GM-CSF enhances inflammatory cytokine expression in a JAK2/STAT5-dependent manner

We next sought to understand how GM-CSF signaling regulates cytokine production. Consistent with our *in vivo* observations, addition of rGM-CSF to isolated MCs infected with *Legionella ex vivo* increased IL-1α, IL-1β, TNF, and IL-12 protein and mRNA levels compared to PBS alone (Figure 6A and S5A). Although *Legionella* infection did not induce GM-CSF production by MCs (Figure S5B), interestingly, BMDMs made GM-CSF in response to *Legionella* (Figure S6A). Critically, this GM-CSF production was important for the ability of BMDMs to produce cytokines, as *Legionella-*infected *Csf2*^*-/-*^ BMDMs produced significantly less IL-1α and IL-1β protein and mRNA compared to WT BMDMs (Figure S6B and S6C). Furthermore, administration of anti-GM-CSF blocking antibodies to *Legionella-*infected BMDMs led to reduced IL-1α and IL-1β protein and mRNA levels, compared to isotype control-treated BMDMs (Figure S6B and S6C). These data indicate that GM-CSF acts directly on myeloid cells to enhance cytokine production.

**Fig. 6.**
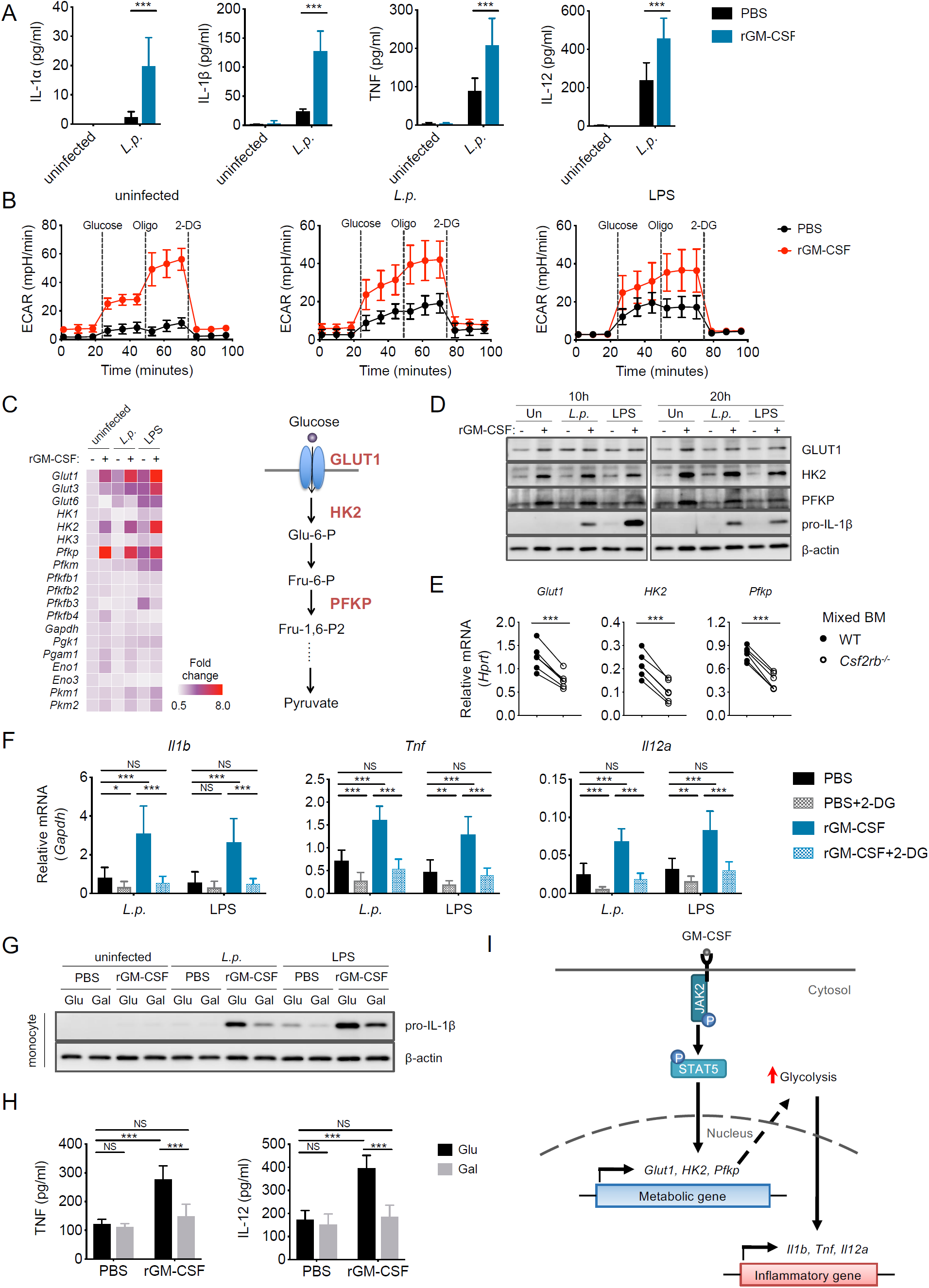
GM-CSF metabolically reprograms monocytes to undergo increased aerobic glycolysis, which enhances inflammatory cytokine production. (A) Isolated WT MCs were uninfected or infected with *L*.*p*. (MOI=5) and given 10 ng/mL rGM-CSF or PBS vehicle control. IL-1α, IL-1β, TNF, and IL-12 levels at 12 hpi are shown. (B) Extracellular acidification rate (ECAR) of uninfected, *L*.*p*.*-*infected (MOI=5), or LPS-treated (10 ng/ml) MCs incubated with rGM-CSF (10 ng/ml) or PBS vehicle control for 12 h before and after sequential treatment with glucose (10mM), oligomycin (Oligo) (1μM), and 2-DG (50 mM). (C) Heatmap depicting the fold-change in transcript levels of genes encoding glucose transporters and glycolytic enzymes in LPS- or *L*.*p*.*-*treated MCs incubated with or without rGM-CSF for 10 h relative to uninfected cells without rGM-CSF. Graphical model depicts an abbreviated version of the glycolytic pathway showing upregulated GLUT1, HK2 and PFKP expression following rGM-CSF treatment. (D) Immunoblot analysis of GLUT1, HK2, PFKP, pro-IL-1β, and β-actin in the lysates of LPS or *L*.*p*.*-* treated MCs incubated with or without rGM-CSF at 10 and 20 hpi. (E) *Glut1, HK2* and *Pfkp* transcript levels in WT and *Csf2rb*^*-/-*^ Ly6C^hi^ MCs from the lungs of 50% WT/50% *Csf2rb*^*-/-*^ →WT mixed BM chimeras at 24 hpi. Each line represents the paired values of WT and *Csf2rb*^*-/-*^ MCs from a given mouse. (F) *Il1b, Tnf*, and *Il12a* transcript levels in LPS- or *L*.*p*.*-*treated MCs treated with 10 mM 2-DG or vehicle control in the presence of rGM-CSF or PBS vehicle control at 6 hpi. (G) Immunoblot analysis of pro-IL-1β and β-actin in the lysates of WT MCs incubated in media containing glucose (10 mM) or galactose (10 mM) that were then uninfected, infected with *L*.*p*. (MOI=5), or treated with LPS (10 ng/ml) in the presence of rGM-CSF or PBS vehicle control for 10 hours.. (H) TNF and IL-12 levels in the supernatants of WT MCs incubated in media containing glucose or galactose, infected with *L*.*p*., and treated with rGM-CSF or PBS vehicle control at 10 hpi. (I) Graphical model depicting that GM-CSF/JAK2/STAT5 signaling upregulates *Glut1, HK2* and *Pfkp* expression and promotes aerobic glycolysis, which is critical for maximal inflammatory cytokine expression. Data shown are the pooled results of three (A, B, F and H) or two (C) independent experiments with triplicate wells per condition in each experiment. Data shown in E are the pooled results of two independent experiments with 3 mice per experiment. Data shown in D and G are representative of two independent experiments. NS, not significant; *p < 0.05, **p < 0.01 and ***p < 0.001 (unpaired t test for A and C; Wilcoxon matched-pairs signed rank test for E; one-way ANOVA with Turkey’s multiple comparisons test for F and H). See also Figure S5, S6, S7, and S8

GM-CSF receptor signaling leads to JAK2 activation and subsequent phosphorylation of the transcription factor STAT5 (*39, 40*). Consistently, we observed a high frequency of phospho-STAT5-positive MCs, AMΦ, and cDCs from the lungs of *Legionella-*infected mice following *ex vivo* GM-CSF treatment (Figure S6D). Similarly, WT BMDMs exhibited robust STAT5 phosphorylation following *Legionella* infection, whereas phosphorylated STAT5 was absent in *Csf2*^*-/-*^ BMDMs, and reduced in a dose-dependent manner by the JAK2 inhibitor NVP-BSK805 (Figure S6E). Notably, both the JAK2 inhibitor NVP-BSK805 and the STAT5 inhibitor Pimozide reduced *ll1a, Il1b, Tnf*, and *Il12a* transcript levels in MCs (Figure S5C and S5D) and *Il1a, Il1b*, and *Ccl2* transcript levels in BMDMs (Figure S6F and S6G) in a dose-dependent manner following rGM-CSF treatment and *Legionella* infection. Thus, GM-CSF-dependent JAK2/STAT5 signaling plays an important role in inducing inflammatory cytokine expression during infection.

### GM-CSF-dependent glycolytic activity is required for optimal inflammatory cytokine production

We next asked how STAT5 might regulate inflammatory gene transcription. One possibility is that STAT5 binds to the promoters or enhancers of inflammatory cytokine genes. Analysis of a previously published chromatin immunoprecipitation sequencing (ChIP-seq) dataset, in which genome-wide STAT5 binding patterns were examined in rGM-CSF-treated DCs (*41*), revealed STAT5 peaks at the 5’ upstream regions of *Il1a* and *Il1b*, in addition to the known GM-CSF-regulated gene *Ccl2* (*42*). In contrast to *Il1a* and *Il1b*, STAT5 was not enriched at *Tnf* and *Il12a* loci (*41*). As *Tnf* and *Il12a* are co-regulated with *Il1a* and *Il1b* by GM-CSF (Figure 6A and S5A), we considered whether alternative mechanisms mediate GM-CSF-dependent expression of inflammatory genes. TLR activation results in increased glycolytic capacity of myeloid cells, and aerobic glycolysis is required for TLR-dependent expression of inflammatory genes (*43-48*). Notably, GM-CSF can enhance LPS-induced glycolysis in BMDMs, and glucose metabolism contributes to GM-CSF-mediated inflammatory gene expression (*49*). We thus hypothesized that in addition to its role in promoting STAT5 binding to a subset of inflammatory genes, GM-CSF promotes glycolytic reprogramming of myeloid cells, and that this enhanced glycolysis contributes to optimal inflammatory gene expression during infection.

We first asked whether GM-CSF causes metabolic alterations in MCs by measuring glycolysis and mitochondrial oxidative phosphorylation through analysis of the extracellular acidification rate (ECAR) and oxygen consumption rate (OCR), respectively. *Legionella* infection or LPS treatment of purified MCs *ex vivo* led to increased glycolysis over uninfected MCs (Figure 6B). Addition of rGM-CSF led to even greater increases in glycolytic activity compared to MCs incubated with *Legionella* or LPS alone (Figure 6B). In contrast, rGM-CSF treatment did not alter mitochondrial respiration in response to *Legionella* or LPS (Figure S7A). Furthermore, the rGM-CSF-mediated increase in glycolysis was blocked by pharmacological inhibition of JAK2 or STAT5 (Figure S7B). These data indicate that in addition to mediating STAT5 enrichment at a subset of inflammatory genes, the GM-CSF/JAK2/STAT5 axis enhances glycolysis.

We next sought to understand how GM-CSF enhances glycolytic activity in MCs, and examined whether GM-CSF upregulates expression of glycolysis-related genes. Indeed, we found that rGM-CSF treatment of LPS- or *Legionella-*infected MCs robustly increased the transcript and protein levels of key rate-limiting glycolytic genes, particularly the glucose transporter *Glut1*, hexokinase *HK2*, and phosphofructokinase *Pfkp*, but not of downstream glycolysis-related genes (Figure 6C and 6D), compared to MCs treated with LPS or *Legionella* alone. Moreover, GM-CSF-mediated upregulation of *Glut1, HK2*, and *Pfkp* was blocked by pharmacological inhibition of JAK2 or STAT5 (Figure S7C). We next investigated whether GM-CSF signaling promotes expression of glycolysis-related genes *in vivo* by examining WT and *Csf2rb*^*-/-*^ MCs from mixed BM chimeras. Intriguingly, *Csf2rb*^*-/-*^ MCs had a significant defect in the expression of *Glut1, HK2, Pfkp* (Figure 6E), as well as enolase *Eno1* (Figure S7D), but not of other glycolysis-related genes, relative to WT MCs within the same mouse during pulmonary *Legionella* infection. These data suggest that GM-CSF reprograms MCs for increased glycolysis by upregulating *Glut1, Hk2*, and *Pfkp* expression, and that this could contribute to GM-CSF-dependent inflammatory cytokine production.

Consistent with this possibility, treatment of MCs with 2-DG, a synthetic glucose analog used to block glycolysis (*50*), significantly suppressed rGM-CSF-dependent induction of *Il1b, Tnf, and Il12a* (Figure 6F). However, 2-DG also impairs oxidative phosphorylation (*51*). To address whether glycolysis is specifically required for GM-CSF-dependent inflammatory gene expression, we employed media containing galactose as a sole carbon source, which must be metabolized by the Leloir pathway before entering glycolysis, resulting in a substantial reduction in glycolytic flux that effectively inhibits glycolysis (*52-54*). Incubation of MCs in galactose-only media suppressed the GM-CSF-mediated increase in glycolysis and did not alter mitochondrial respiration, compared to MCs in glucose-only media (Figure S8A and S8B). Moreover, LPS- or *Legionella-*treated MCs incubated in galactose-only media had a substantial defect in pro-IL-1β, TNF, or IL-12 production following rGM-CSF treatment compared to MCs in glucose-only media (Figure 6G and 6H). These data indicate that GM-CSF-induced glycolysis is critical for optimal cytokine production by MCs in response to infection. Collectively, these results indicate that GM-CSF-dependent glycolysis and transcriptional regulation are integrated to promote optimal inflammatory cytokine production in MCs, thus promoting successful control of pulmonary infection.

## Discussion

Within the lung, alveolar macrophages serve not only as initial sentinels of infection, but also as potential reservoirs for replication by many pulmonary pathogens. DAMPs such as IL-1, that are released from infected alveolar macrophages, have an important role as a contingency system that promotes inflammation in response to pathogens that block PRR signaling (*18, 55*). Here, we have found that the initially infected macrophages are insufficient on their own to direct a subsequent robust inflammatory response by bystander myeloid cells, as IL-1 released by infected macrophages does not directly license bystander myeloid cells to produce inflammatory cytokines. Rather, we find that the alveolar epithelium is critical for amplifying immune responses by serving as a key signaling relay between infected alveolar macrophages and recruited myeloid cells. We found that IL-1 instructs alveolar epithelial cells to produce GM-CSF, which metabolically reprograms bystander myeloid cells to undergo increased glycolysis in order to support optimal inflammatory cytokine production. GM-CSF enforced additional production of IL-1α and IL-1β by myeloid cells, indicating that GM-CSF and IL-1 participate in a feedforward loop. Furthermore, GM-CSF was required for maximal TNF and IL-12 production, and subsequent IFNγ production, cytokines that are required to control *Legionella* infection (*56*).

Our findings define a three-way cell circuit initiated by release of DAMPs from infected macrophages that is relayed by the lung epithelium to license myeloid cells for maximal cytokine production. Why such a three-way communication network is necessary for optimal responses against *Legionella* is unclear. However, each alveolus is patrolled by a surprisingly low number of alveolar macrophages, with approximately one alveolar macrophage for every three alveoli (*3, 4*), indicating that a signal amplification system is needed to generate appropriate responses. AECII, which comprise 60% of alveolar epithelial cells, provide a perfectly poised system to amplify the initial alarm sounded by infected macrophages in the lung. We propose that this crosstalk between the alveolar epithelium and myeloid cells allows for the robust amplification of inflammatory responses, particularly against immunoevasive pathogens.

We found that GM-CSF reprogrammed MCs to undergo increased aerobic glycolysis, which was required for maximal cytokine production. Although GM-CSF was sufficient to drive glycolysis, it was not sufficient on its own for cytokine production by purified monocytes *ex vivo*, as bacterial infection or LPS stimulation was required, suggesting that PRR sensing of PAMPs is involved. PRRs, such as TLRs, lead to NF-κB signaling and inflammatory gene expression (*57*). TLR signaling also results in glycolytic reprogramming of myeloid cells that is critical for inflammatory gene expression, in part through stabilizing hypoxia-inducible factor-1α, to transcriptionally regulate *Il1b* expression, and by supporting the de novo synthesis of fatty acids to allow for ER and Golgi expansion and accommodate increased protein production (*44-48*). Thus, GM-CSF signaling likely cooperates with PRR signaling for maximal glycolytic reprogramming to drive cytokine production. Further studies are needed to determine how GM-CSF, PRR signaling, and metabolic reprogramming collaborate for maximal inflammatory gene expression.

GM-CSF can be produced by a variety of hematopoietic and stromal cell types, including lymphocytes, myeloid cells, epithelial cells, and endothelial cells (*58-60*). Although we found that macrophages make GM-CSF in response to *Legionella* under *in vitro* conditions, in the context of *in vivo* infection, we found that AECII are the major producers of GM-CSF in response to IL-1. Bronchial epithelial cells can also produce GM-CSF in response to IL-1 during allergic airway responses (*61*). Whether other tissues use similar mechanisms to mediate stromal GM-CSF production is unknown. In the steady-state gut, RORγt^+^ innate lymphoid cells (ILCs) are the major source of GM-CSF in response to IL-1β released by macrophages upon sensing the microbiota (*60*). Colonic epithelial cells produce GM-CSF in response to DSS-induced injury to promote repair of the colonic mucosa (*62*), and in response to invasive bacteria *in vitro* (*63*). Epithelial cells at other barrier tissue sites could also be an important source of GM-CSF by sensing IL-1 or other DAMPs released in response to invasive pathogens.

It is becoming increasingly appreciated that the alveolar epithelium regulates essential immune functions during infection, such as the production of chemokines that recruit immune cells to the lung (*30, 38, 64*). Our findings indicate that the alveolar epithelium also directly modifies the function of recruited myeloid cells by reprogramming their metabolism and licensing them to produce inflammatory cytokines important for anti-bacterial defense within the lung. Understanding other signals sensed by the lung epithelium to enhance myeloid cell function may provide insight into how to modulate local inflammatory responses within the lung for the purposes of treating pulmonary infection.

## Methods

### Mice

*Il1α*^*-/-*^ and *Il1β*^*-/-*^ (*65*), *Il1r1*^*-/-*^ (*66*), *Csf2*^*-/-*^ (*34*), *Csf2rb*^*-/-*^ (*67*), *Il1r1*^*fl/fl*^ ;Spc-cre-ERT2 and *Il1r1*^*r/r*^ ;Spc-cre-ERT2 (generated by crossing *Il1r1*^*fl/fl*^ (*68*) or *Il1r1*^*r/r*^ (*69*) and Spc-cre-ERT2 (*70*) mice in our facility), and *Il1r1*^*fl/*fl^ ;CD11c-cre mice (generated by crossing *Il1r1*^*fl/fl*^ and CD11c-cre (*71*) mice in our facility) were bred and housed under specific-pathogen free conditions at the University of Pennsylvania. Wild-type C57BL/6J or B6.SJL controls or littermate controls were either purchased from Jackson Laboratories or bred in house. Mice were used at 8-10 weeks of age, gender-matched, and littermates of the same sex were randomly assigned to experimental groups. All animal studies were performed in compliance with the federal regulations set forth in the Animal Welfare Act (AWA), the recommendations in the Guide for the Care and Use of Laboratory Animals of the National Institutes of Health, and the guidelines of the University of Pennsylvania Institutional Animal Use and Care Committee. All protocols used in this study were approved by the Institutional Animal Care and Use Committee at the University of Pennsylvania (protocol #804928).

### Cell lines and primary cell cultures

For generating bone marrow-derived macrophages, bone marrow cells from gender- and age-matched wild-type C57BL/6J mice and *Csf2*^*-/-*^ mice were differentiated in RPMI supplemented with 20% FBS, 30% L929 cell supernatant, and 1% penicillin/streptomycin solution (P/S) at 37°C in a 5% CO2 incubator for 7 days. Macrophages were replated in RPMI with 10% FBS and 15% L929 cell supernatant without P/S. Primary murine monocytes were isolated from C57BL/6J bone marrow using the Monocyte Isolation Kit (Miltenyi Biotec, 130-100-629) according to the manufacturer’s instructions. Monocytes were replated in RPMI with 10% FBS without P/S. For experiments involving media containing glucose or galactose, monocytes were replated in RPMI with 10% FBS for 4 hours, and then the media was replaced with RPMI supplemented with 10% dialyzed FBS lacking glucose prior to infection and stimulation.

To isolate primary murine AECII, the lungs of gender- and age-matched WT C57BL/6J mice and *Il1r1*^*-/-*^ mice (6-8 weeks) were perfused via the right ventricle with 5 ml DPBS, lavaged with 1 ml DPBS three times, inflated with a 1.5 ml mixture of 1 ml low melting agarose (3% w/v) and 500 μl 4U/ml dispase (Worthington Biochem, LS02104), and incubated for 1 h at room temperature (RT). The lung lobes were gently grinded on 100 μm mesh soaked in DMEM, and then filtered through 40 μm mesh. The cell suspension was centrifuged at 300 g for 10 min. The cell pellet was resuspended in FACS buffer containing 5% FBS. AECII were then purified by fluorescence-activated cell sorting (see Method Details). AECII cells were plated into a Matrigel (Corning, 354230) pre-coated TC plate and cultured with DMEM supplemented with charcoal-stripped 5% FBS, 10 ng/ml keratinocyte growth factor (PeproTech, 100-19), 10 μM Rock inhibitor (Selleck Chemicals, S1049), and 1% P/S at 37°C in a 5% CO2 incubator for the first 2 days, and then replaced with the same media but without Rock inhibitor for the next 4 to 5 days.

### Bacterial cultures

*Legionella pneumophila* JR32-derived flagellin-deficient Δ*flaA* mutant (*72, 73*) and Lp02-derived (thymidine auxotroph) Δ*flaA* mutant (*72*) were cultured on charcoal yeast extract (CYE) agar plates for 48 h at 37°C prior to infection. JR32 Δ*flaA* strain was used for all in vivo infection and for measuring intracellular bacterial replication within BMDMs. Lp02 Δ*flaA* strain was used for all other in vitro experiments.

### Bone marrow chimeric mice experiments

Wild-type B6.SJL mice (CD45.1 background) or knockout (KO) mice (*Il1r1*^*-/-*^, *Csf2*^*-/-*^, and *Csf2rb*^*-/-*^, CD45.2 background) received a lethal dose of irradiation (1096 Rads). 6 hours later, mice were injected retro-orbitally with freshly isolated WT, KO or mixed (1:1 ratio) bone marrow cells (5×10^6^ cells per mouse). All chimeras were provided antibiotic-containing water (40 mg trimethoprim and 200 mg sulfamethoxazole per 500 ml drinking water) for four weeks after irradiation and subsequently provided acidified water without antibiotics for another 4 or 8 weeks. The reconstitution of hematopoietic cells (proportion of donor CD45 cells among total CD45 cells) in the lung was analyzed by flow cytometry.

### Infection of mice

Mice were anesthetized by intraperitoneal injection of ketamine (100mg/kg) and xylazine (10 mg/kg) solution and then infected intranasally with *Legionella pneumophila* JR32 Δ*flaA* strain (10^6^ CFU per mouse). At the indicated time points, the bronchoalveolar lavage fluid (BALF) was harvested with 1 ml cold PBS. The lung lobes were excised and digested with dissociation solution (20 U/ml DNAseI, 240 U/ml Collagenase IV, and 5% FBS in PBS) at 37°C for 30 min. Mechanical dissociation of the lung tissue was performed using the Miltenyi GentleMACS™ Dissociator. The lung homogenates were incubated with red blood cell lysis buffer on ice for 5 min and then quenched with cold PBS. The cell pellet was resuspended with FACS buffer to generate a single cell suspension and then filtered through nylon mesh prior to flow cytometric analysis. To measure bacterial CFUs within the lung, lung lobes were excised and placed into in 5 ml sterile water, homogenized using the Miltenyi GentleMACS™ Dissociator, and cell lysates were then plated onto CYE agar plates. Bacterial loads were determined by calculating the number of bacterial colony forming units (CFU) per gram of lung (CFU/g).

### Antibody-based neutralization of GM-CSF in vivo

C57BL/6J mice were IP injected with 250 μg of anti-GM-CSF antibodies (Bio-X-Cell, clone MP1-22E9) or 250 μg of isotype control antibodies (Bio-X-Cell, clone 2A3) 16 hours prior to infection, followed by a second injection when mice were infected. For experiments in which bacterial loads in the lungs were measured, mice were also IP injected at days 1 and 2 post-infection with 125 μg of anti-GM-CSF or isotype control antibodies.

### Cytokine production assays

For in vivo experiments, the first 1 ml of bronchoalveolar lavage was used for cytokine measurements. For in vitro experiments, tissue culture plates containing cells were centrifuged 5 min at 300 g, and then supernatants were collected for analysis. Most cytokines were measured using commercial mouse ELISA kits. IL-12 was measured by ELISA using purified anti-mouse IL-12 p40/70 antibody (BD Biosciences, 551219) and biotin rat anti-mouse IL-12 p40/70 antibody (BD Biosciences, 554476). The Luminex assay was performed using the MILLIPLEX™ Mouse Cytokine/Chemokine Magnetic Bead Panel (Millipore-Sigma) with technical assistance from the Human Immunology Core at the University of Pennsylvania Perelman School of Medicine.

### Flow cytometry

To analyze cell populations in the BALF and lung, cell suspensions were treated with fluorescent Zombie Yellow™ dye (live/dead stain; Biolegend) for 15 min at RT. Anti-CD16/CD32 antibody was used to block Fc receptors. Fluorescently conjugated antibodies, including anti-CD45 (BioLegend, clone 30-F11), anti-CD45.1 (BioLegend, clone A20), anti-CD45.2 (BioLegend, clone 104), anti-Ly6G (BioLegend, clone 1A8), anti-Ly6C (BioLegend, clone HK1.4), anti-Siglec F, anti-CD11c (BioLegend, clone N418), anti-CD11b (BioLegend, clone M1/70), anti-MHC II (BioLegend, clone M5/114.15.2), anti-CD31 (BioLegend, clone MEC13.3), anti-EpCAM (Thermo Fisher Scientific, clone G8.8), anti-T1a (BioLegend, clone 8.1.1), anti-Sca-1(BD Biosciences, clone D7), and anti-CD49f (BioLegend, clone GoH3), were used for surface staining. For intracellular cytokine staining, cell suspensions were incubated with Brefeldin A (0.1%) and monensin (0.066%) solution for 4 hours at 37°C prior to all staining, treated with BD Cytofix/Cytoperm™ buffer for 20 min at 4°C, and stained with antibodies specific for TNFα (Thermo Fisher Scientific, clone MP6-XT22), IL-12 (BioLegend, clone C15.6), GM-CSF (BioLegend, clone MP1-31G6), IL-1α (Thermo Fisher Scientific, clone ALF-161), or IL-1β (Thermo Fisher Scientific, clone NJTEN3) in perm/wash buffer for 30 min at 4°C. Isotype antibodies were used as negative controls. To detect STAT5 phosphorylation, cells were fixed with BD PhosFlow™ Lyse/Fix Buffer for 10 min at 37°C, permeabilized by BD PhosFlow™ Perm Buffer III for 30 min on ice, and incubated with PE-conjugated anti-phospho-STAT5 (Y694) antibody (BD Biosciences, clone 47). For reactive oxygen species (ROS) measurements, cell suspensions were incubated with CellROX™ Deep Red reagent (Thermo Fisher Scientific, C10491) at a final concentration of 5 μM at for 30 min at 37°C, then immediately analyzed on the BD LSR II flow cytometer. Data were analyzed with FlowJo software (version 10.3)

### Fluorescence-activated cell sorting of lung cell subsets

Single cell lung suspensions were incubated with Zombie Yellow™ dye, anti-CD16/CD32 antibody, and fluorescently-labeled antibodies (see Key Resource Table). Non-hematopoietic cell subsets were identified as follows: alveolar type II epithelial cell (AECII, CD45-CD31-EpCAM+T1a-CD49f^low^), alveolar type I epithelial cell (AECI, CD45-CD31-EpCAM+T1a+CD49f^low^), endothelial cell (EnC, CD45-CD31+EpCAM-), bronchial alveolar stem cell (BASC, CD45-CD31-EpCAM^hi^ CD49f^hi^ Sca-1+). Hematopoietic cell subsets were identified as follows: neutrophil (NΦ, CD45+Ly6G+CD11b+), alveolar macrophage (AMΦ, CD45+SiglecF+CD11c+CD11b-), monocyte (MC, CD45+Ly6G-SiglecF-CD11c-CD11b+Ly6C^high^), conventional dendritic cell (cDC, CD45+Ly6G-SiglecF-MHCII^hi^ CD11c^hi^), NK cell (CD45+CD3-NK1.1+), NKT cell (CD45+CD3+NK1.1+), αβT cell (CD45+CD3+TCRβ+), and γδT cell (CD45+CD3+TCR γ/δ+). Individual cell types were isolated using a BD FACSAria™ Cell Sorter.

### RNA isolation and quantitative RT-PCR

Total RNA was isolated using RNeasy Mini Kit (Qiagen, 74106). cDNA was generated using SuperScript II Reverse Transcriptase. Primers (see Key Resource Table) were synthesized by Integrated DNA Technologies, Inc. (Coralville, IA). Quantitative RT-PCR was performed using SsoFast™ EvaGreen® Supermix with Low ROX (Bio-Rad, 1725212) on a CFX96™ Real-Time PCR Detection System (Bio-Rad). Primers used are listed in Table 1. mRNA expression of each gene relative to *Gapdh* or *Hprt* was calculated using the formula 2^−ΔCt^.

**Table 1.**
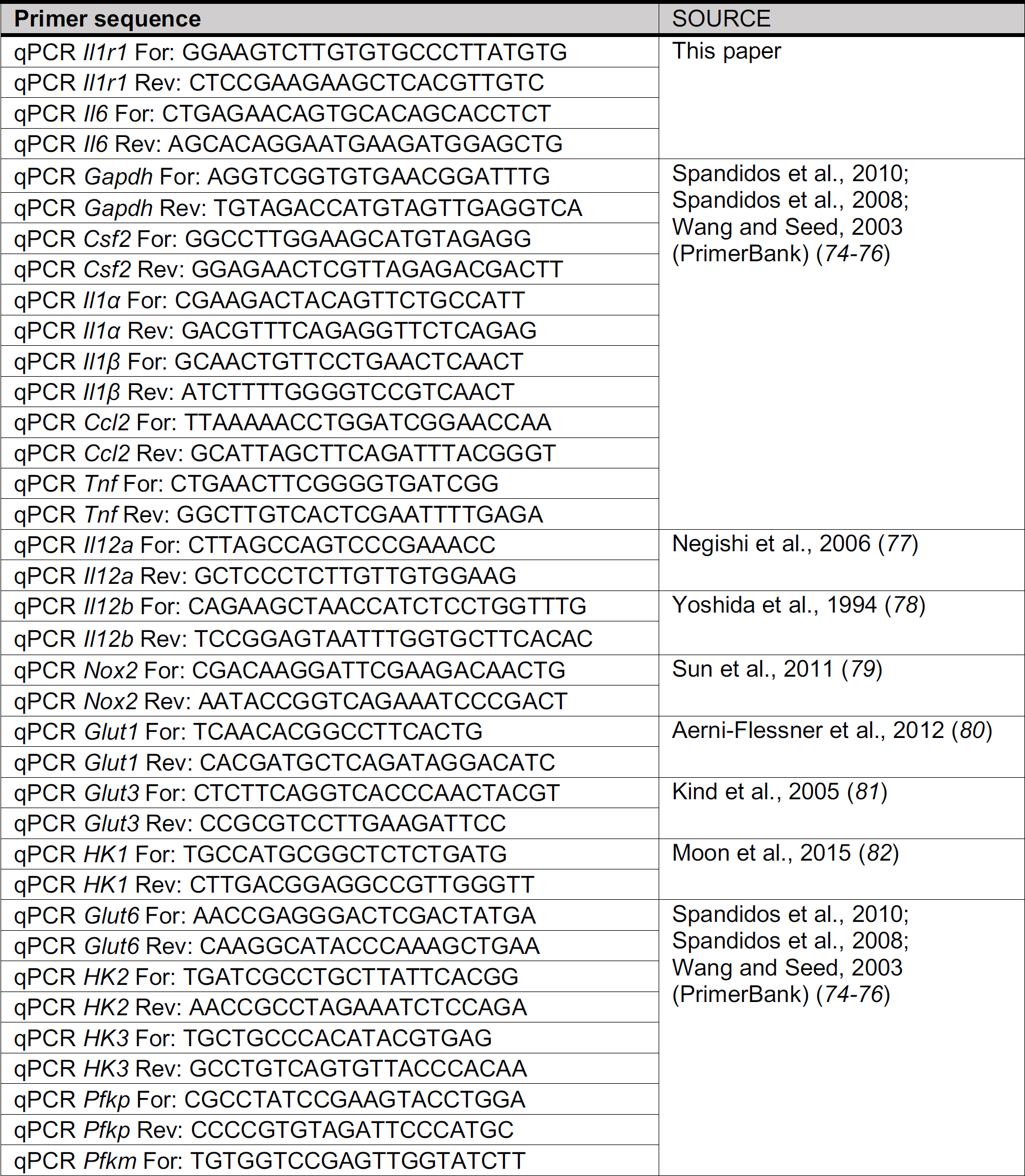
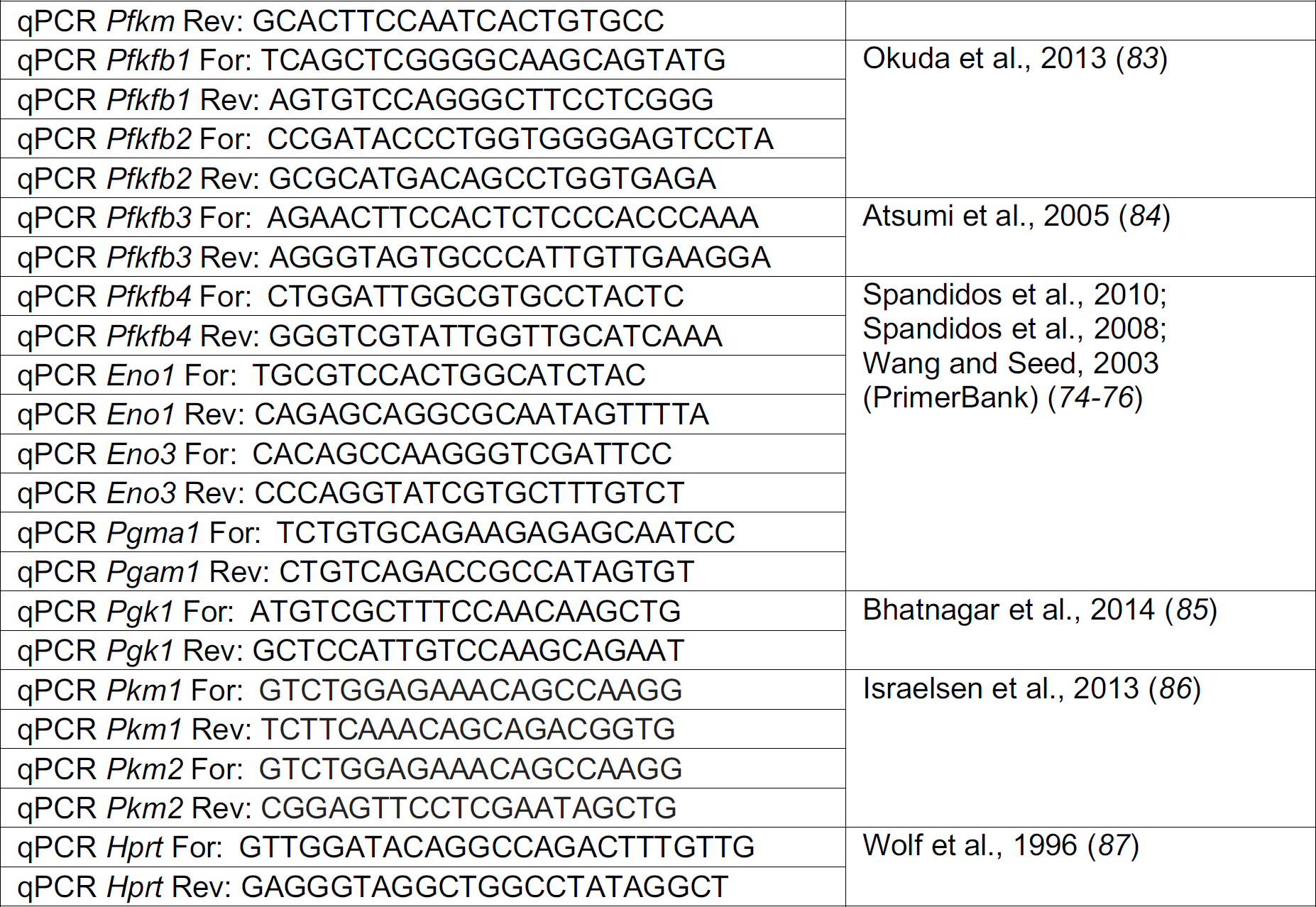
List of primers used for qPCR.

### Immunoblot analysis

Cells were lysed in 1x SDS-PAGE loading buffer supplemented with 1x Halt™ Protease and Phosphatase Inhibitor Cocktail. Protein samples were separated by SDS-PAGE and then transferred to PVDF membranes. Primary antibodies specific for pro-IL-1β (1:1000, CST, 12242), STAT5 (1:1000, CST, 9363), phospho-STAT5 (Y694) (1:1000, CST, 9351), GLUT1 (1:1000, Thermo Fisher, PA1-46152), HK2 (1:1000, CST, 2867), PFKP (1:1000, Thermo Fisher, PA5-28673) and β-actin (1:1000, CST, 4967) were used. Anti-rabbit or mouse HRP-linked IgG antibody was used as a secondary antibody (1:2000, CST, 7074 or 7076). Detection was performed using SuperSignalTM Substrates (Thermo Fisher Scientific, 34076 and 34095).

### Glycolysis and oxidative phosphorylation analysis

Primary mouse bone marrow monocytes were obtained as described above and seeded into Seahorse XF96 cell culture microplates (10^5^ cells per well) containing RPMI media with 10% FBS. Four hours later, the cells were infected with or without Lp02 Δ*flaA* strain (MOI=5), or stimulated with LPS (from E.coli, 10 ng/ml), followed by treatment with rGM-CSF or PBS vehicle control. After 12 hours of stimulation, the media was replaced with fresh Seahorse XF base media supplemented with 2 mM L-glutamine for the glycolysis test, or additionally supplemented with 10 mM glucose and 1 mM pyruvate for the oxidative phosphorylation test, and pH was adjusted to 7.4. Plates were incubated without CO_2_ for 45 min in a 37°C incubator and then loaded into the Seahorse XFe96 analyzer. The Seahorse XF Glycolysis Stress Test Kit (Agilent Technology, 103020-100) and Cell Mito Stress Test Kit (Agilent Technology, 103015-100) were used to measure the extracellular acidification rate (ECAR, mpH/min) and oxygen consumption rate (OCR, pmol/min), respectively. Glucose (10 mM), Oligomycin (1 µM) and 2-DG (50 mM) for ECAR measurements, and Oligomycin (1 µM), FCCP (1 µM) and Rot/AA (0.5 µM) for OCR measurements were injected at indicated time points. ECAR and OCR were calculated using Wave software V2.4.

### Statistical analysis

All statistical analyses were performed using GraphPad Prism software (GraphPad). Data are shown as mean or mean ± SD. Statistical significance was determined by performing t tests for two unpaired groups, Wilcoxon matched-pairs signed rank test for paired two groups, or one-way ANOVA with Tukey’s multiple comparisons test for three or more groups as indicated in each figure legend. *P* values less than 0.05 were considered to be statistically significant.

## Acknowledgements

We thank Igor Brodsky for scientific advice, discussion, and critical reading of the manuscript; members of the Brodsky and Shin labs, Christopher Hunter, and Terri Laufer for insightful discussions and scientific advice; the Pancreatic Islet Cell Biology Core of the Institute for Diabetes, Obesity, and Metabolism for assistance with Seahorse assays; the Flow Cytometry and Cell Sorting Resource Laboratory for flow cytometric technical support; and Yoichiro Iwakura for generously providing *Il1α*^*-/-*^ and *Il1β*^*-/-*^ mice.

## Funding

This work was supported by NIH/NIAID grants R01AI118861 (SS) and R01AI123243 (SS), a Linda Pechenik Montague Investigator Award from the University of Pennsylvania Perelman School of Medicine (SS), and a Burroughs-Wellcome Fund Investigators in the Pathogenesis of Infectious Diseases Award (SS).

## Author Contributions

X.L., A.M.H, and S.S. conceived and designed experiments. X.L., M.A.B., and A.M.H. performed experiments. X.L., M.A.B., A.M.H., and S.S. analyzed and interpreted data. X.L. prepared figures. X.L. and S.S. wrote the paper.

## Competing interests

Authors declare no competing interests.

## Data and Materials Availability

All data is available in the main text or supplementary materials.

## Supplementary Materials

Figures S1-S8

