## Supplemental Figures for "Infected macrophages engage alveolar epithelium to metabolically reprogram myeloid cells and promote antibacterial inflammation"

Figure S1

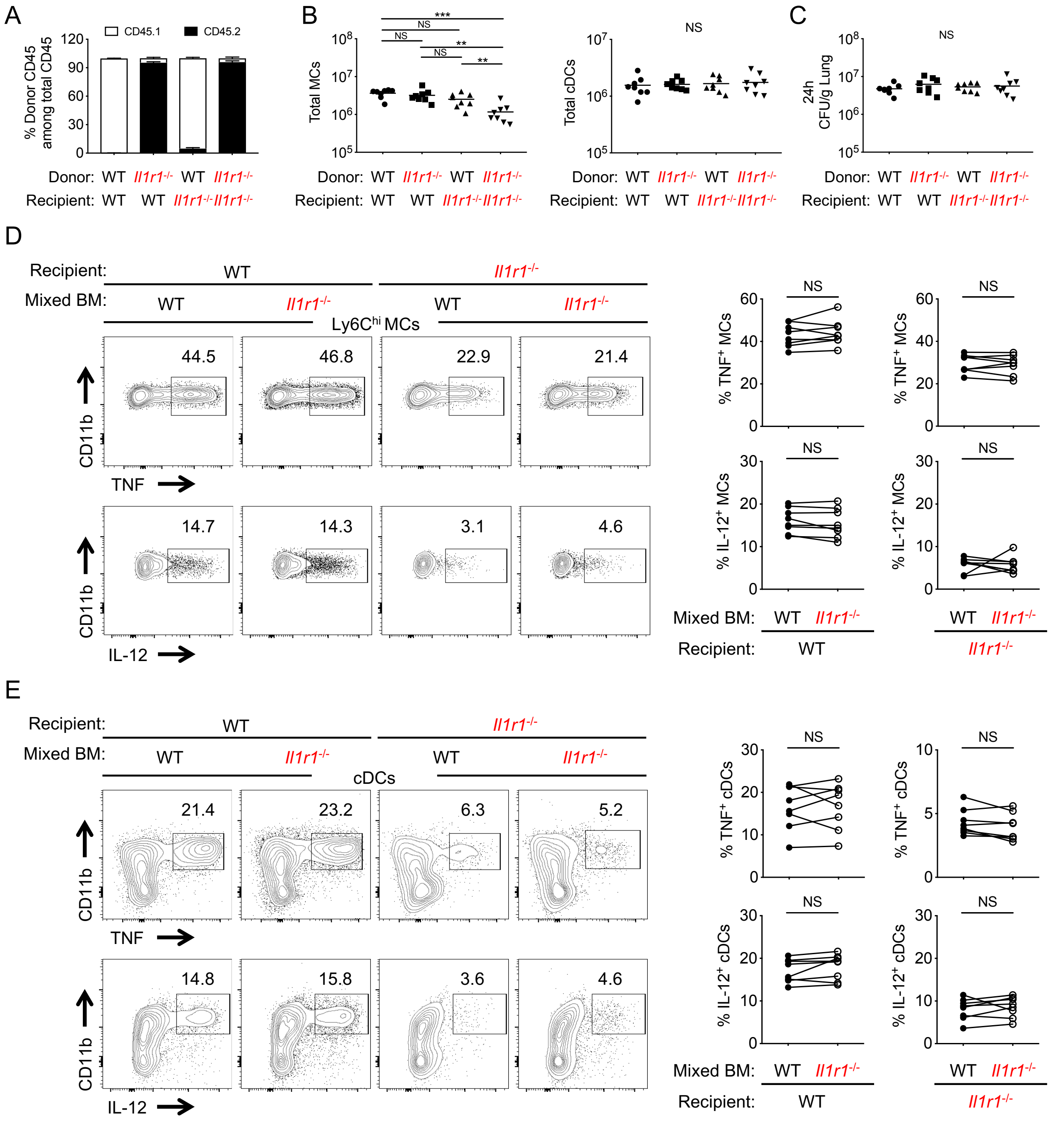


**Figure S1. Cell-extrinsic IL-1R signaling mediates TNF and IL-12 production by Ly6C$^{hi}$ MCs and cDCs, related to Figure 1**

(A) Graphs depicting the percentage of donor CD45$^{+}$ cells among total CD45$^{+}$ cells in the lungs of BM chimeric mice 12 weeks after BM transplantation (mean ± SD).

(B) Graphs depicting the total numbers of MCs and cDCs in the lungs of BM chimeric mice at 24 hpi.

(C) *L.p.* CFUs in the lungs of chimeric mice at 24 hpi.

(D and E) Representative flow cytometric plots showing intracellular staining for TNF and IL-12 in Ly6C$^{hi}$ MCs (D) and cDCs (E) from the lungs of 50% WT/50% *Il1r1*$^{-/-}$ →WT or 50% WT/50% *Il1r1*$^{-/-}$ →*Il1r1*$^{-/-}$ mixed BM chimeras at 24 hpi. Quantification of the frequency of TNF- or IL-12-producing WT or *Il1r1*$^{-/-}$ Ly6C$^{hi}$ MCs (D) and cDCs (E) are shown. Each line represents the paired values of WT and *Il1r1*$^{-/-}$ cells from a given mouse.

Data shown are the pooled results of two independent experiments with 4 mice per group in each experiment. NS, not significant; $**p < 0.01$; and $***p < 0.001$ (one-way ANOVA with Turkey's multiple comparisons test for B and C; Wilcoxon matched-pairs signed rank test for D and E).

Figure S2

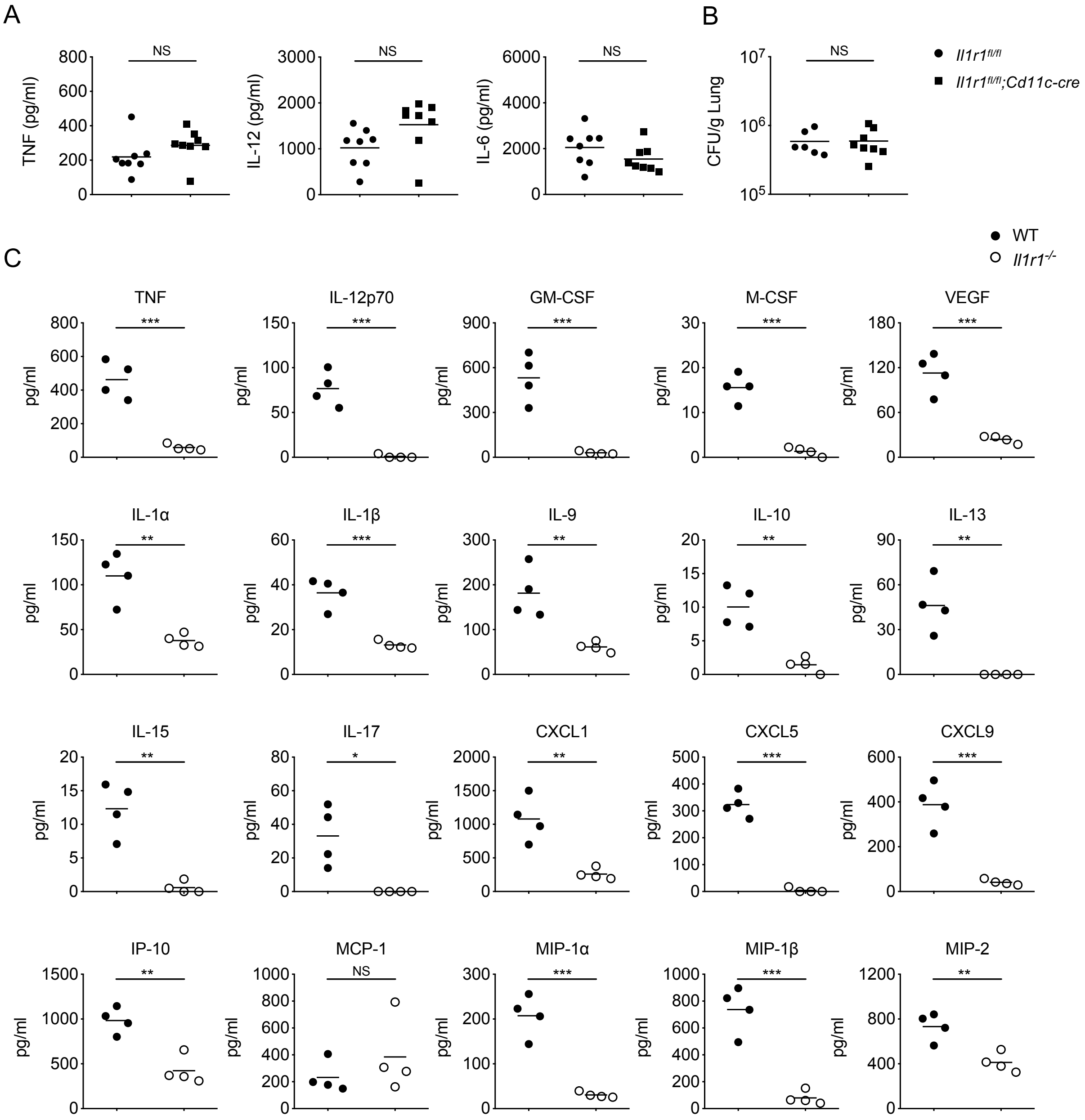


**Figure S2. IL-1R expression on CD11c-expressing cells is dispensable for cytokine production and bacterial clearance, and mice lacking IL-1R have a defect in the production of multiple cytokines and chemokines, related to Figures 2 and 3**

(A-B) *Il1r1$^{fl/fl}$;Cd11c-cre* mice and *Il1r1$^{fl/fl}$* littermates were intranasally infected with *L.p.* TNF, IL-12, and IL-6 levels in the BAL at 24 hpi (A). *L.p.* CFUs in the lungs of mice at 72 hpi (B). Data shown are the pooled results of two independent experiments with 3-4 mice per group in each experiment. NS, not significant (unpaired t test).

(C) Cytokine and chemokine levels in the BAL of WT and *Il1r1*$^{-/-}$ mice at 24 hpi were measured by Luminex assay. Data shown are the pooled results of 4 mice per group. NS, not significant; $*p<0.05$; $**p < 0.01$; and $***p < 0.001$ (unpaired t test)

### Figure S3

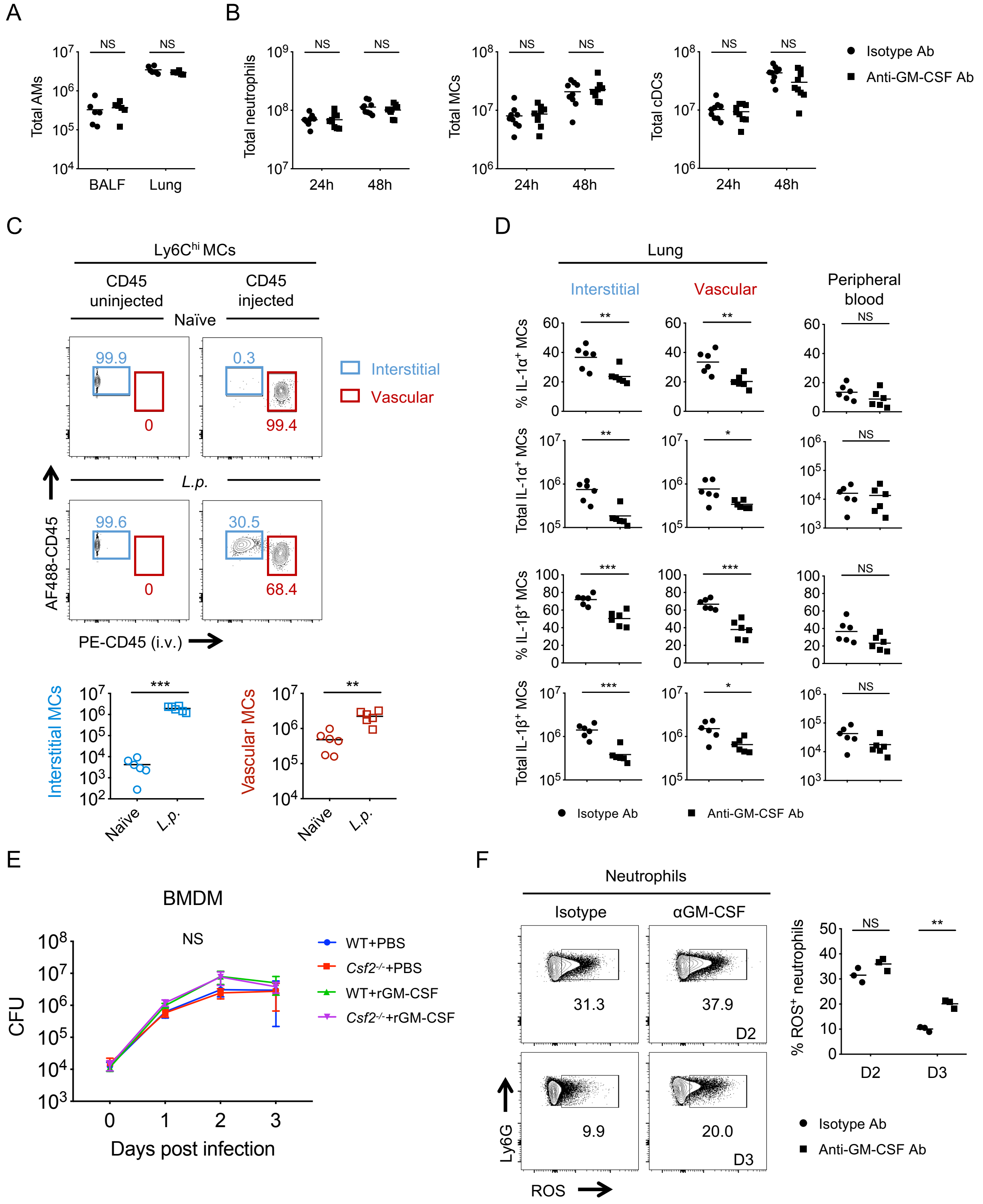

A
Total AMs
NS
NS
BALF
Lung
B
Total neutrophils
Total MCs
Total cDCs
NS
24h
48h
Isotype Ab
Anti-GM-CSF Ab
C
Ly6Chi MCs
CD45 uninjected
CD45 injected
Naïve
L.p.
99.9
0
0.3
99.4
99.6
0
30.5
68.4
Interstitial
Vascular
AF488-CD45
PE-CD45 (i.v.)
Interstitial MCs
Vascular MCs
***
**
Naïve
L.p.
D
Lung
Interstitial
Vascular
Peripheral blood
% IL-1α+ MCs
Total IL-1α+ MCs
% IL-1β+ MCs
Total IL-1β+ MCs
**
*
***
NS
Isotype Ab
Anti-GM-CSF Ab
E
BMDM
NS
CFU
WT+PBS
Csf2-/-+PBS
WT+rGM-CSF
Csf2-/-+rGM-CSF
Days post infection
F
Neutrophils
Isotype
αGM-CSF
31.3
37.9
D2
9.9
20.0
D3
Ly6G
ROS
% ROS+ neutrophils
NS
**
D2
D3
Isotype Ab
Anti-GM-CSF Ab

**Figure S3. GM-CSF is critical for local inflammatory cytokine production by myeloid cells within the lung, related to Figure 4**

(A) C57BL/6 mice were intraperitoneally injected with anti-GM-CSF or isotype control Ab. 16 hours later, total numbers of alveolar macrophages in the BALF and lung were quantified.

(B) Total numbers of neutrophils, Ly6C$^{hi}$ MCs, and cDCs in the lungs of *L.p.*-infected mice following anti-GM-CSF or isotype Ab injection were quantified at 24 and 48 hpi.

(C) *L.p.*-infected or naïve WT mice were either uninjected or intravenously injected with PE-labeled anti-CD45 antibodies three minutes prior to harvesting of lung tissue. Representative flow cytometric plots showing the percentages of interstitial MCs (AF488-CD45$^{+}$, PE-CD45$^{-}$) and vascular MCs (AF488-CD45$^{+}$, PE-CD45$^{+}$) in the lungs of infected and uninfected mice at 24 hpi. The total number of interstitial and vascular MCs in the lungs of infected and naïve mice were quantified at 24 hpi.

(D) Percentage and total number of IL-1$\alpha$- or IL-1$\beta$-producing Ly6C$^{hi}$ MCs from the interstitial or vascular lung compartments or the peripheral blood of infected mice administered anti-GM-CSF or isotype Ab at 24 hpi.

(E) Intracellular bacterial replication was assessed in WT and *Csf2*$^{-/-}$ BMDM following treatment with or without 10ng/ml recombinant GM-CSF protein (rGM-CSF) at days 0, 1, 2 and 3 post-infection (MOI=1).

(F) Representative flow cytometric plots and graph depicting the percentage of reactive oxygen species (ROS)-positive neutrophils in the BAL of anti-GM-CSF or isotype Ab-treated mice at days 2 or 3 post-infection.

Data shown are the pooled results of two (A, C, and D) or three (B) independent experiments with 3 mice. Data shown in E are the pooled results of two independent experiments with triplicate wells per condition for each experiment. Data shown in F represent 3 mice per group. NS, not significant; $*p<0.05$, $**p < 0.01$ and $***p < 0.001$ (unpaired t test for A-D and F; one-way ANOVA with Turkey's multiple comparisons test for E).

Figure S4

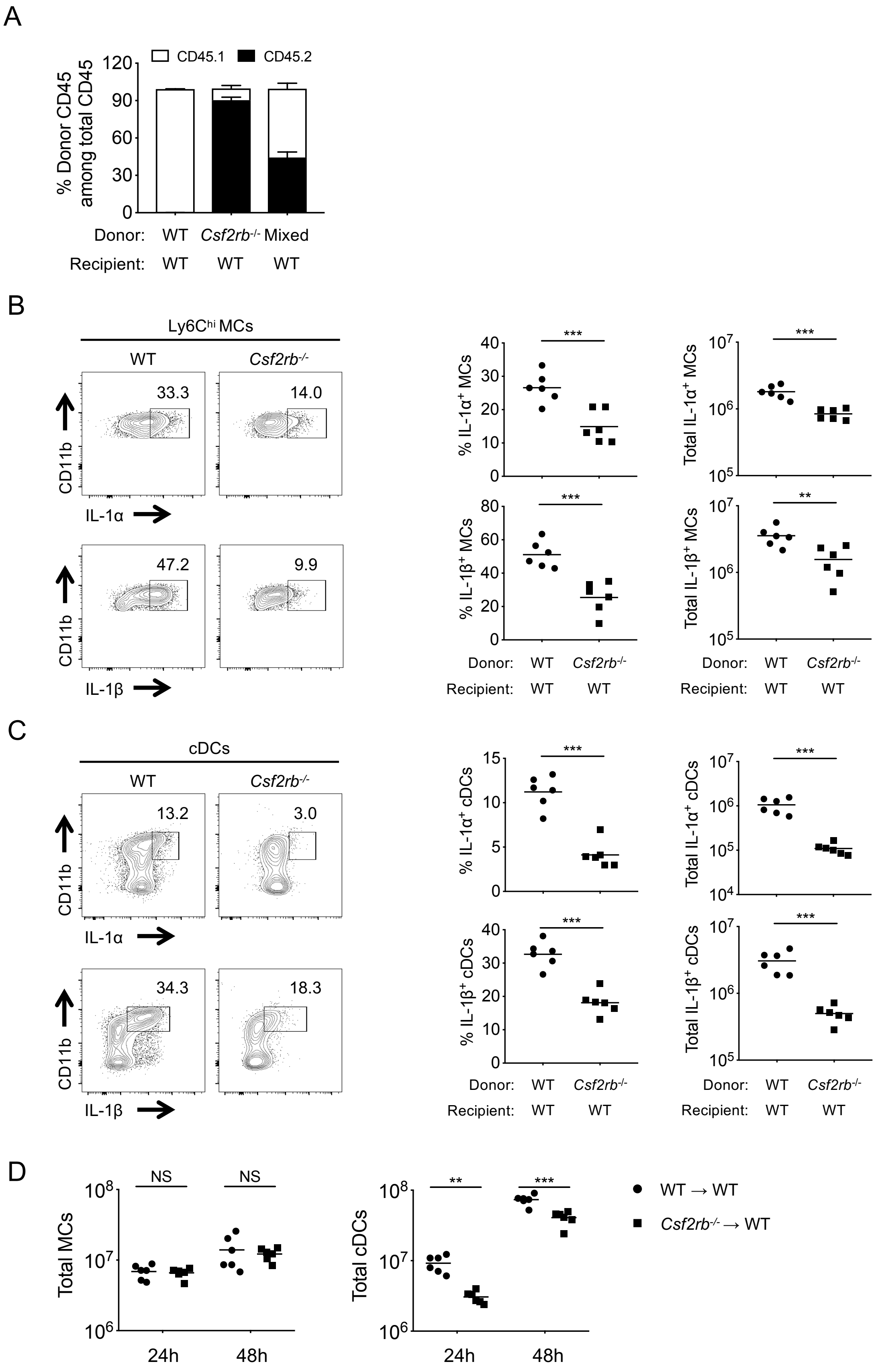
A
CD45.1
CD45.2
% Donor CD45 among total CD45
120
90
60
30
0
Donor: WT Csf2rb-/- Mixed
Recipient: WT WT WT
B
Ly6Chi MCs
WT
Csf2rb-/-
33.3
14.0
47.2
9.9
CD11b
IL-1α
IL-1β
% IL-1α+ MCs
Total IL-1α+ MCs
% IL-1β+ MCs
Total IL-1β+ MCs
***
**
Donor: WT Csf2rb-/-
Recipient: WT WT
C
cDCs
13.2
3.0
34.3
18.3
% IL-1α+ cDCs
Total IL-1α+ cDCs
% IL-1β+ cDCs
Total IL-1β+ cDCs
D
Total MCs
Total cDCs
NS
24h
48h
WT → WT
Csf2rb-/- → WT

**Figure S4. GM-CSF receptor expression in the hematopoietic compartment is required for inflammatory cytokine production following infection, related to Figure 5**

(A) Graphs depicting the percentage of donor CD45 cells among total CD45 cells in the lungs of BM chimeric mice 8 weeks after BM transplantation (mean ± SD).

(B and C) Representative flow cytometric plots showing intracellular staining for IL-1$\alpha$ and IL-1$\beta$ in Ly6C$^{hi}$ MCs (B) and cDCs (C) from the lungs of chimeric WT→WT and *Csf2rb*$^{-/-}$→WT mice at 24 hpi. Graphs depict the frequency and total numbers of IL-1$\alpha$- and IL-1$\beta$-producing Ly6C$^{hi}$ MCs (B) and cDCs (C) at 24 hpi, as determined by intracellular cytokine staining and flow cytometry.

(D) Total numbers of Ly6C$^{hi}$ MCs and cDCs in the lungs of chimeric WT→WT and *Csf2rb*$^{-/-}$→WT mice at 24 and 48 hpi.

Data shown are the pooled results of two independent experiments with 3 mice per chimera in each experiment. NS, not significant; $**p < 0.01$ and $***p < 0.001$ (unpaired t test).

Figure S5

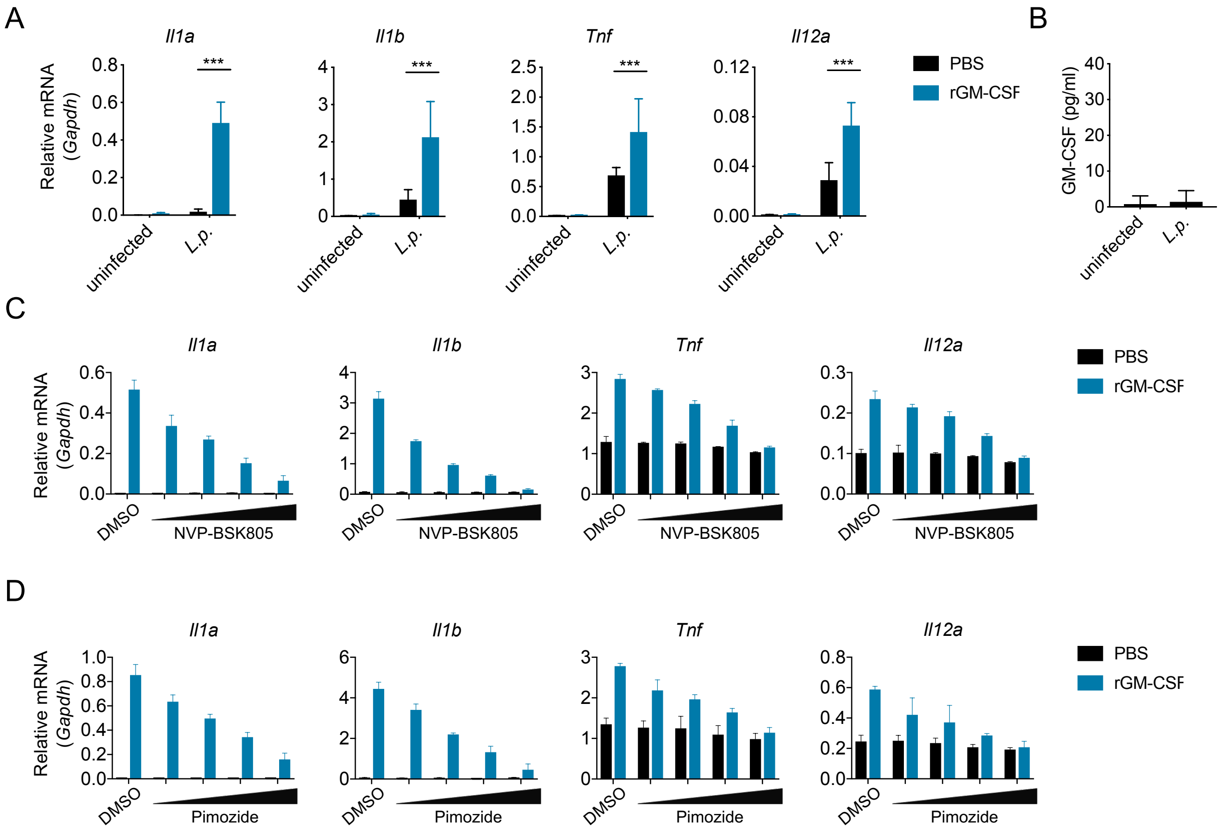


**Figure S5. GM-CSF-dependent JAK2/STAT5 signaling is required for inflammatory cytokine expression by monocytes, related to Figure 6**

(A) *Il1a*, *Il1b*, *Tnf,* and *Il12a* transcript levels in *L.p.*-infected MCs supplemented with 10 ng/mL rGM-CSF or PBS vehicle control at 12 hpi.

(B) GM-CSF levels in the supernatants of MCs that were uninfected or infected with *L.p.* (MOI=5) at 12 hpi.

(C and D) *Il1a*, *Il1b*, *Tnf,* and *Il12a* transcript levels in MCs infected with *L.p.* (MOI=5), supplemented with 10 ng/mL rGM-CSF or PBS vehicle control, and treated with JAK2 inhibitor NVP-BSK805 (250, 500, 1000, or 2000 nM) (C), STAT5 inhibitor Pimozide (1, 2, 2.5, or 5 $\mu$M) (D), or DMSO vehicle control, at 12 hpi.

Data shown are the pooled results of three (A and B) or two (C and D) independent experiments. NS, not significant; ***$p < 0.001$ (unpaired t test for A, C, and D).

### Figure S6

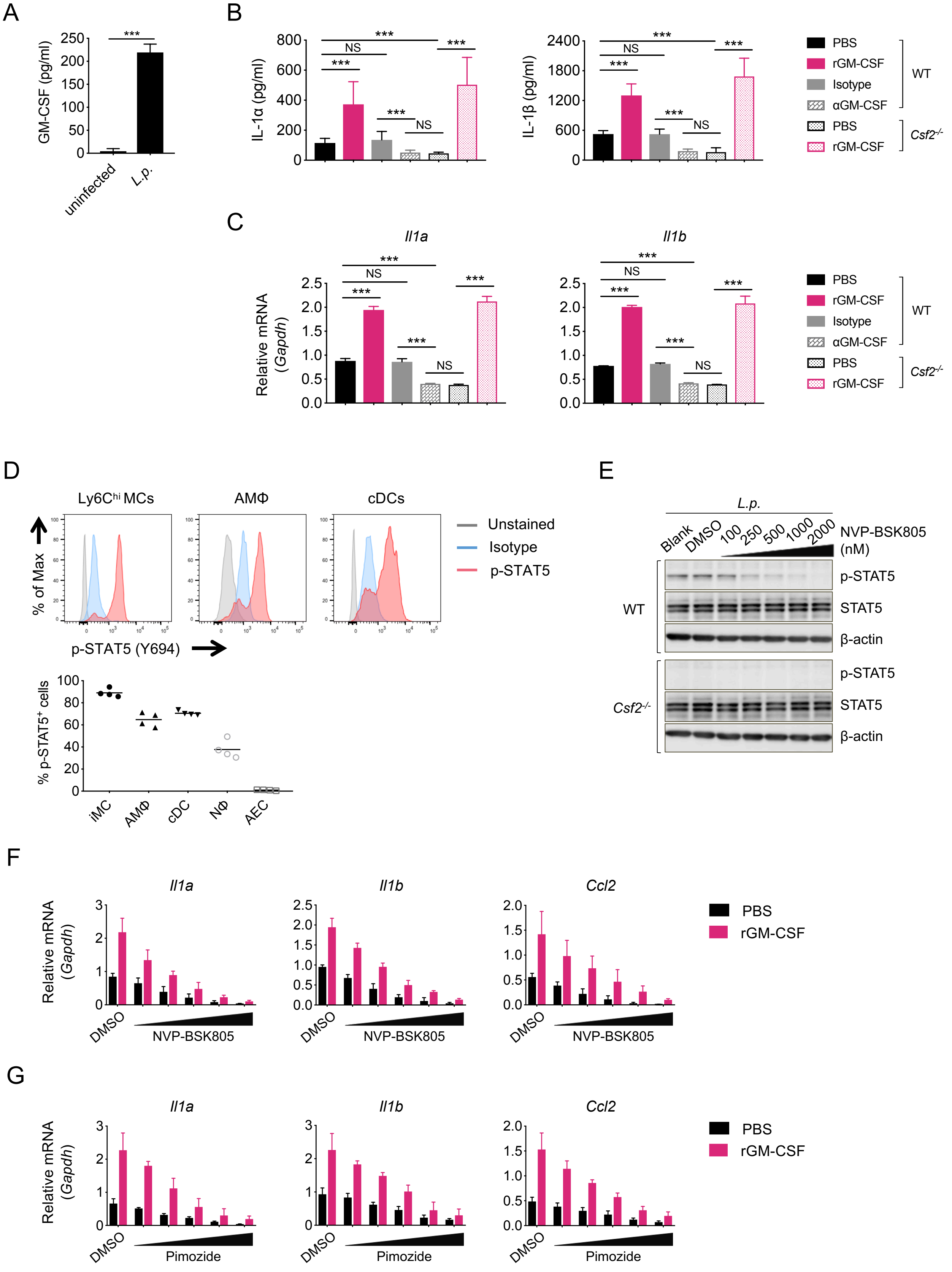

A
GM-CSF (pg/ml)
uninfected
L.p.
B
IL-1α (pg/ml)
IL-1β (pg/ml)
NS
PBS
rGM-CSF
Isotype
αGM-CSF
PBS
rGM-CSF
WT
Csf2-/-
C
Il1a
Il1b
Relative mRNA (Gapdh)
D
Ly6Chi MCs
AMΦ
cDCs
% of Max
p-STAT5 (Y694)
Unstained
Isotype
p-STAT5
% p-STAT5+ cells
iMC
AMΦ
cDC
NΦ
AEC
E
L.p.
Blank
DMSO
100
250
500
1000
2000
NVP-BSK805 (nM)
p-STAT5
STAT5
β-actin
WT
Csf2-/-
F
Il1a
Il1b
Ccl2
DMSO
NVP-BSK805
G
Pimozide

**Figure S6. GM-CSF-dependent JAK2/STAT5 signaling is required for maximal inflammatory cytokine expression in macrophages, related to Figure 6**

(A) GM-CSF levels in the supernatants of BM-derived macrophages (BMDMs) that were uninfected or infected with *L.p.* (MOI=5) at 12 hpi.

(B and C) IL-1$\alpha$ and IL-1$\beta$ supernatant levels (B) and transcriptional levels of *Il1a* and *Il1b* (C) in WT and *Csf2*$^{-/-}$ BMDMs infected with *L.p.* (MOI=5) and supplemented with 10 ng/mL rGM-CSF or PBS, or 10 µg/ml anti-GM-CSF or isotype Abs.

(D) Representative flow cytometry histogram showing levels of phosphorylated-STAT5 in Ly6C$^{hi}$ MCs, AMΦ, and cDCs isolated from the lungs of infected WT mice at 24 hpi and stimulated *in vitro* with rGM-CSF (10 ng/ml) at 37°C for 15 min. Depicted are unstained cells (grey line), cells stained with isotype antibodies (blue line), or cells stained with anti-pSTAT5 (Y694) antibodies (red line). Graph shows the percentage of p-STAT5-positive cells in the indicated cell types.

(E) Immunoblot analysis of phosphorylated and total STAT5 and $\beta$-actin in the lysates of WT and *Csf2*$^{-/-}$ BMDMs infected with *L.p.* (MOI=5) and treated with indicated concentrations of JAK2 inhibitor NVP-BSK805 or DMSO vehicle control at 12 hpi.

(F and G) *Il1a*, *Il1b*, and *Ccl2* transcript levels in WT BMDMs infected with *L.p.* (MOI=5), supplemented with 10 ng/mL rGM-CSF or PBS, and treated with JAK2 inhibitor NVP-BSK805 (100, 250, 500, 1000, or 2000 nM) (G), or STAT5 inhibitor Pimozide (0.5, 1, 2, 2.5, or 5 $\mu$M) (H), or DMSO vehicle control, at 12 hpi.

Data shown in A-C are the pooled results of three independent experiments. Data shown in D and E are representative of two independent experiments. Data shown in F and G are the pooled results of two independent experiments. NS, not significant; $***p < 0.001$ (unpaired t test for A, F, and G; one-way ANOVA with Turkey's multiple comparisons test for B and C).

### Figure S7

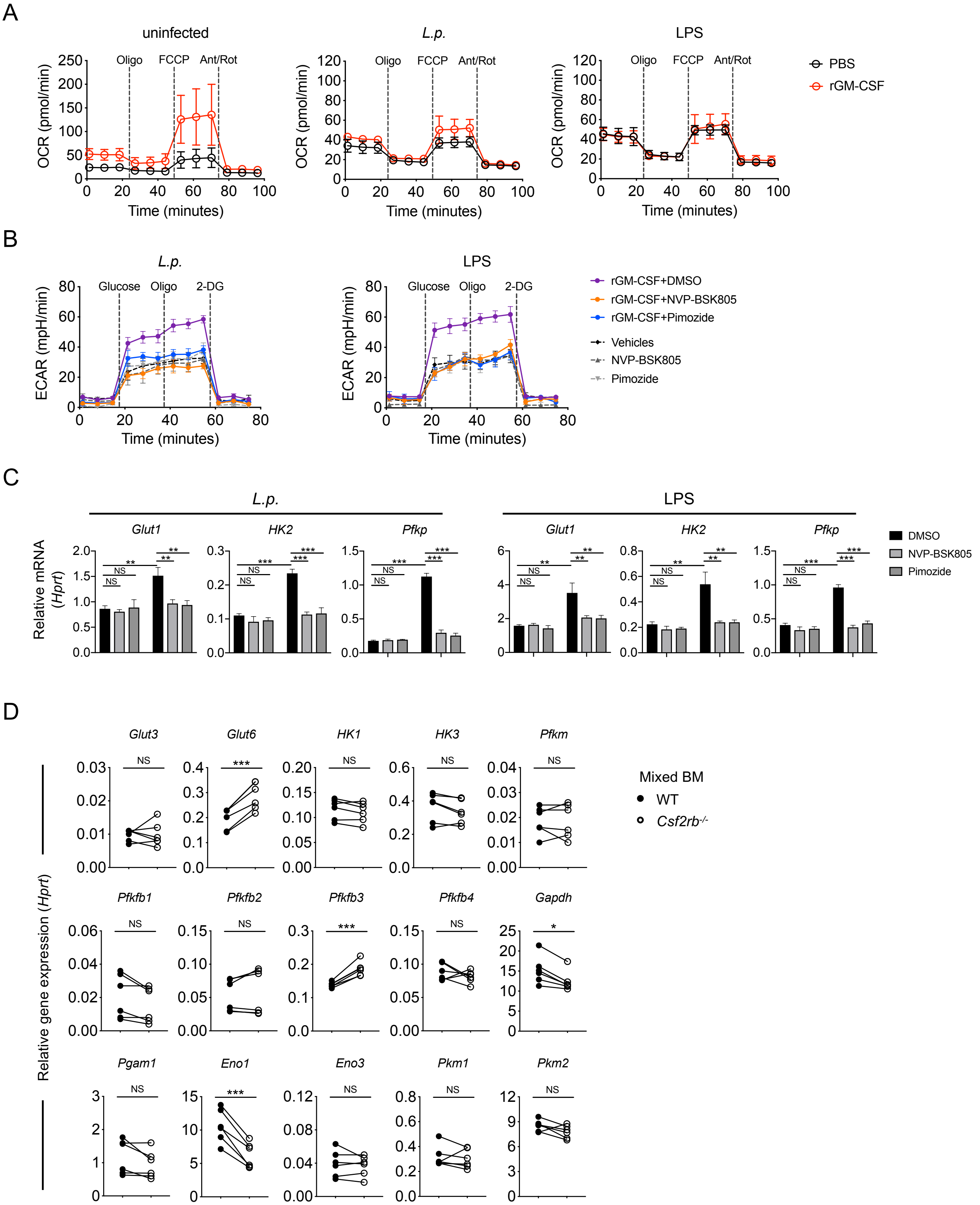
A
uninfected
L.p.
LPS
OCR (pmol/min)
Oligo
FCCP
Ant/Rot
Time (minutes)
PBS
rGM-CSF
B
ECAR (mpH/min)
Glucose
2-DG
rGM-CSF+DMSO
rGM-CSF+NVP-BSK805
rGM-CSF+Pimozide
Vehicles
NVP-BSK805
Pimozide
C
Relative mRNA (Hprt)
Glut1
HK2
Pfkp
DMSO
NS
D
Glut3
Glut6
HK1
HK3
Pfkm
Pfkfb1
Pfkfb2
Pfkfb3
Pfkfb4
Gapdh
Pgam1
Eno1
Eno3
Pkm1
Pkm2
Relative gene expression (Hprt)
Mixed BM
WT
Csf2rb-/-

**Figure S7. GM-CSF-dependent JAK2/STAT5 signaling promotes aerobic glycolysis and glycolytic gene expression, related to Figure 6**

(A) Oxygen consumption rate (OCR) of monocytes that were uninfected, infected with *L.p.* (MOI=5), or treated with LPS (10 ng/ml) in the presence of rGM-CSF (10 ng/ml) or PBS vehicle control for 12 hours, before and after sequential treatment with oligomycin (Oligo) (1μM), carbonyl cyanide-4-(trifluoromethoxy) phenylhydrazone (FCCP) (1μM), and antimycin/rotenone (Ant/Rot) (0.5μM).

(B and C) MCs were pretreated with JAK2 inhibitor NVP-BSK805 (2μM) or STAT5 inhibitor pimozide (5 μM) or DMSO for 30 mins, then treated with rGM-CSF or PBS and treated with *L.p.* or LPS. ECAR was measured 12 hpi (B), or *Glut1*, *HK2* and *Pfkp* transcript levels relative to *Hprt* were measured at 10 hpi (C).

(D) Transcript levels of glycolysis-related genes in WT and *Csf2rb*$^{-/-}$ Ly6C$^{hi}$ MCs isolated from the lungs of *L.p.*-infected 50% WT/50% *Csf2rb*$^{-/-}$→WT mixed BM chimeras at 24 hpi. Each line represents paired values of WT and *Csf2rb*$^{-/-}$ MCs from a given mouse.

Data shown are the pooled results of three (A) or two (B and C) independent experiments. Data shown in D are the pooled results of two independent experiments with 3 mice per experiment. NS, not significant; $*p < 0.05$; $**p < 0.01$; and $***p < 0.001$ (unpaired t test for A-C; Wilcoxon matched-pairs signed rank test for D).

Figure S8

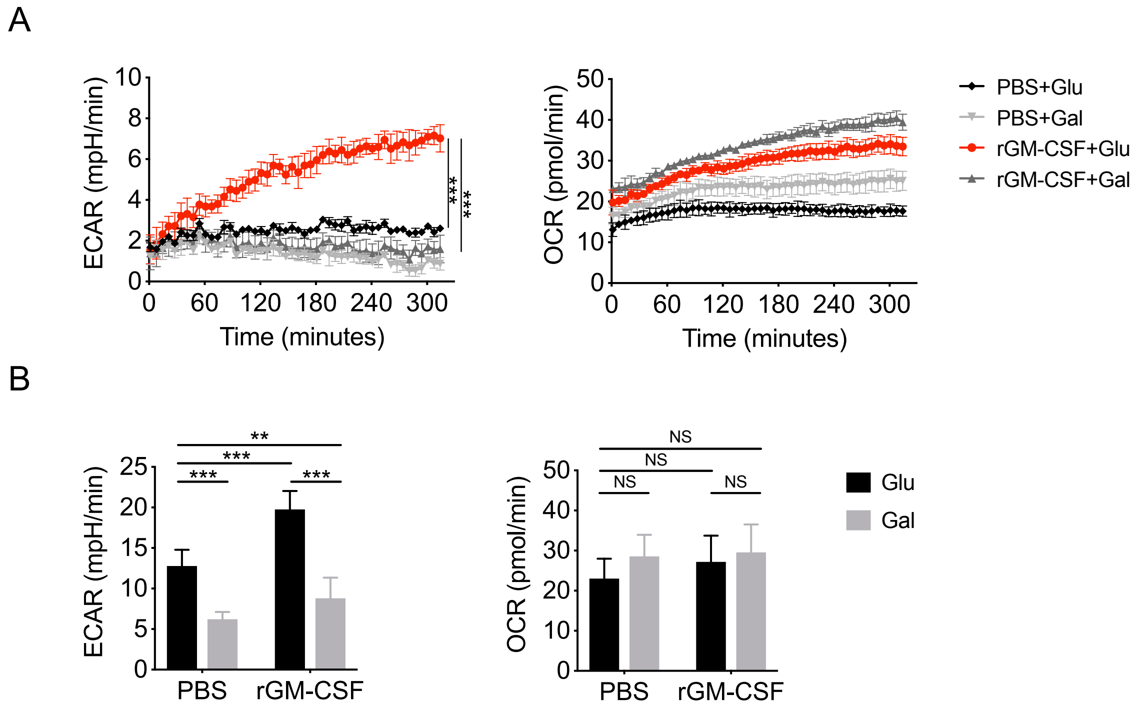


**Figure S8. Galactose-containing media suppresses glycolytic activity but not mitochondrial respiration, related to Figure 6**

(A and B) Real-time ECAR and OCR in monocytes with media containing glucose or galactose, infected with *L.p.,* and treated with rGM-CSF or PBS vehicle control (A). ECAR and OCR at the end-time point for each group are shown (B).

Data shown in A are representative of three independent experiments. Data shown in B are the pooled results of three independent experiments. NS, not significant; ***p < 0.001 (one-way ANOVA with Turkey's multiple comparisons test).
